# Plumage convergence resulting from social mimicry in birds? A tetrachromatic view

**DOI:** 10.1101/2020.03.30.016113

**Authors:** María Alejandra Meneses-Giorgi, Carlos Daniel Cadena

## Abstract

Social mimicry may lead to convergent evolution when interactions with conspecific and heterospecific individuals drive evolution towards similar phenotypes in different species. Several hypotheses accounting for convergence in communication signals based on mechanisms of social mimicry exist, but evaluations of how similar species are given the visual system of receptors of such signals have been ostensibly missing from tests of such hypotheses. We used plumage reflectance measurements and models of avian color discrimination to evaluate the efficacy of visual deception and therefore the plausibility of mimicry hypotheses accounting for plumage convergence among six species of passerine birds in the flycatcher family (Tyrannidae) with strikingly similar plumage. We rejected interspecific social mimicry hypotheses as an explanation for the similarity between one putative model species and putative mimics because deception seems unlikely given the visual system of passerines. However, plumage similarity was consistent with a role for selective pressures exerted by predators because dorsal coloration of putative model and mimic species was indistinguishable by visually oriented raptors. Experiments and behavioral observations are necessary to better characterize social interactions and to test predictions of alternative mimicry hypotheses proposed to account for convergence.

## INTRODUCTION

Convergent evolution, the process through which two or more distinct lineages independently acquire similar traits, reveals that the paths of evolution are not infinite, but may instead be rather restricted. Convergence may happen rapidly or over the course of millions of years by random drift [1] or, more likely, because a given phenotypic trait is repeatedly favored by natural selection in a particular environment [2,3]. Likewise, convergence may also occur via biases in the production of phenotypic variation, such as shared developmental constraints [3–5]. One well-studied form of convergent evolution is mimicry, in which one species (the mimic) evolves to resemble another species (the model), often to deceive a third species (the receptor; [6]).

There are numerous examples of phenotypic convergence among birds [7–15], and several authors have proposed hypotheses to explain this phenomenon in the context of mimicry [7,13,16–22].

Among leading ideas suggested to account for phenotypic convergence in birds, the social mimicry hypothesis [18] posits that convergent similarity in traits like coloration and plumage patterns may evolve to promote efficient communication maintaining cohesion both among conspecifics and heterospecifics in mixed-species flocks. A variant of this hypothesis suggests that rather than maintaining cohesion of mixed flocks, social mimicry serves mainly as an antipredatory adaptation because predation eliminates conspicuous or atypical individuals from populations, thereby promoting phenotypic uniformity [16]. How atypical an animal is in this context must be examined relative to the background [23]; if a predator considers a whole mixed-species flock as the background, then any phenotype forming a distinct minority within it may be a preferred prey, resulting in a selective pressure favoring homogeneity [24]. Consequently, the efficacy of social mimicry to reduce predation [16] depends on the extent to which predators may perceive mixed flocks as homogeneous, which ultimately relies on discrimination abilities determined by their visual system.

An alternative explanation for mimicry not focusing on predation but still considering social interactions suggests that mimicry may serve two purposes: (1) mimics may escape attacks from model species of larger body size, and (2) mimics may deceive species of smaller size and scare them off without further effort [20]. Along the same lines, Prum & Samuelson [19] proposed the Interspecific Social Dominance Mimicry (ISDM) hypothesis, which posits that, given interference competition, smaller species evolve to mimic larger, ecologically dominant competitors to deceive them and thereby avoid attacks. For this mechanism to be plausible, individuals of the model species must confuse individuals of the mimic species as if they were conspecifics based on visual cues like shape, color and plumage patterns despite differences in body size [7,13,19,21]. Therefore, the efficacy of this form of mimicry critically depends on the visual system of model species.

Assessing the plausibility of various mimicry hypotheses has been limited by the lack of explicit tests of the effectiveness of visual deception under models of avian vision (but see [8,9]). Like humans, birds have visual pigments enabling them to acquire information from short (using the *s* cone type), medium (using the *m* cone type), and long wavelengths (using the *l* cone type), but they can also acquire information from ultraviolet and violet wavelengths with an additional pigment (using the *u* or *v* cone type, respectively). Additionally, each of the avian pigments is paired with a pigmented oil droplet type, which is hypothesized to result in better spectral discrimination relative to other vertebrates [25]. The ability to distinguish colors is thought to vary among birds, however, with a pronounced difference in the absorbance peak of the ultraviolet-sensitive (UVS-type) cones present in Passeriformes and Psittaciformes, and the violet-sensitive (VS-type) cones present in all other non-passerines including raptors [26]. Thus, a crucial question one must answer to gauge support for alternative mimicry hypotheses is whether phenotypic similarities between species perceived by humans are sufficient to potentially deceive birds including predators, competitors, and putative models given properties of their visual systems.

We used plumage reflectance measurements of six species of tyrant flycatchers (Passeriformes, Tyrannidae) with strikingly similar plumage patterns to evaluate the efficacy of visual deception and therefore the plausibility of alternative mimicry hypotheses potentially accounting for phenotypic convergence. The species we studied are part of a hypothetical mimicry complex posited to be an example of ISDM consisting of up to two model species of large body size and several putatively mimic species of smaller size [7]. We took reflectance measurements of eight plumage patches and compared plumage coloration for each pair of hypothetical models and mimics, both from the perspective of predatory raptors (using a standard VS vision model) and of models, mimics, and smaller competitors (using a standard UVS vision model) to evaluate the plausibility of deception of different receptors. Because raptors are likely the main diurnal predators of passerine birds [27–30] and detect them by sight [24,31,32], the social mimicry hypothesis that species converge phenotypically to deceive predators [16] predicts that species of flycatchers involved in the mimicry complex should be similar to each other or indistinguishable under the raptor (VS) visual model. Such similarity should be particularly evident in dorsal coloration, under the assumption that predators primarily detect and attack potential prey from above [23,33]. On the other hand, hypotheses positing that species evolve to deceive heterospecifics with which they may compete for resources [7,19,20] predict that tyrant flycatcher species involved in the mimicry complex should have indistinguishable plumage coloration under the passerine (UVS) visual model.

## METHODS

### Study system

We studied six phylogenetically dispersed species in the tyrant-flycatcher family (Figure 1): Boat-billed Flycatcher (*Megarynchus pitangua*, mean body mass 73.5 g, mean size 23 cm) and Great Kiskadee (*Pitangus sulphuratus*, 63.8 g, 22 cm) as hypothetical models, and Lesser Kiskadee (*Pitangus lictor*, 25.5 g, 18 cm), White-bearded Flycatcher (*Phelpsia inornata*, 29.4 g, 17.5cm), Social Flycatcher (*Myiozetetes similis*, 28 g, 16.5 cm), and Rusty-margined Flycatcher (*Myiozetetes cayanensis*, 25.9 g, 16.5 cm) as hypothetical mimics [7,34–36]. All these species show strikingly similar plumage patterns which we refer hereafter to as “kiskadee-like”: black facial mask, white throat, bright yellow underparts, brownish upperparts, and rufous-edged tail and wings (Illustrations in Figure 1; [35,37]). These are all lowland species (mostly ranging from 500m to 1700m) with wide distributional ranges (maps in Figure 1) except for *P. inornata*, which is restricted to the *llanos* of Colombia and Venezuela [37]. The distributional ranges of putative models and mimics overlap extensively, and species generally share habitats in semi-open areas. Despite having overall similar plumage patterns and coloration to the human eye, the species differ in details of the coloration of the head, back, rump and primary feathers, as well as in size and shape of the bill, morphology, and songs [35].

**Figure 1.**
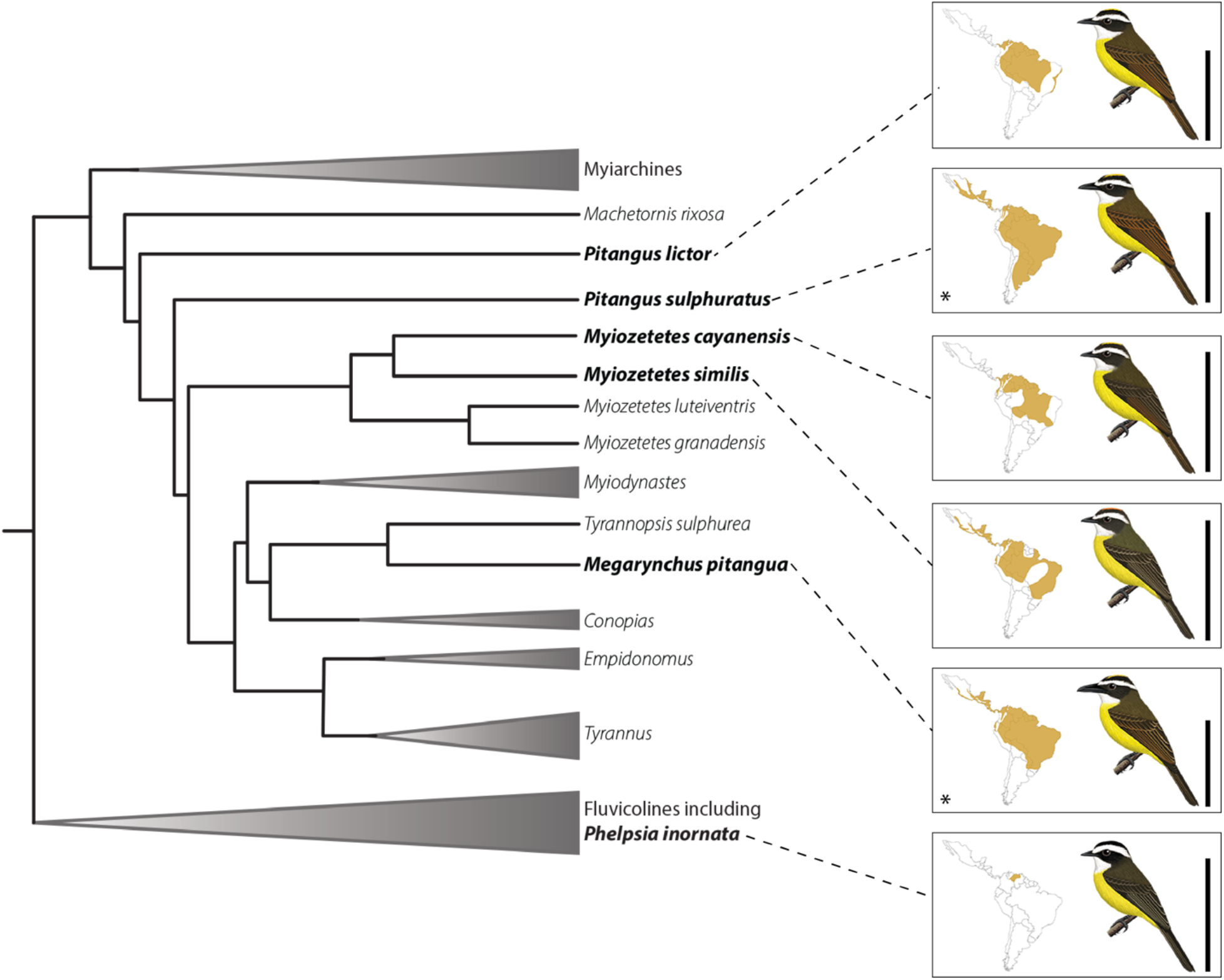
Phylogenetic position of species with kiskadee-like plumage in flycatcher phylogeny illustrates convergence in phenotype. Study species are shown in bold and connected by dotted lines to maps showing their overlapping distributions and illustrations of their strikingly similar plumages. Bars next to illustrations represent 15cm and are scaled to indicate body size of each species. Hypothetical models are indicated with an asterisk. The tree is a schematic based on ongoing analyses of suboscine phylogeny based on thousands of loci in the nuclear genome (M. Harvey et al. in review). Illustrations by Ayerbe-Quiñones [36] reproduced with permission from the author.

### Quantifying plumage similarity

#### Reflectance measurements

We quantified plumage similarity among hypothetical models and mimics using spectrophotometric data obtained from museum specimens from Colombia deposited in the Museo de Historia Natural de la Universidad de los Andes (ANDES), Instituto de Ciencias Naturales de la Universidad

Nacional (ICN), and Instituto de Investigación de Recursos Biológicos Alexander von Humboldt (IAvH). We took reflectance measurements using an Ocean Optics USB4000 spectrophotometer and a DH-2000 deuterium halogen light source coupled with a QP400-2-UV-VIS optic fiber with a 400 μm diameter. We measured reflectance of eight plumage patches: crown, back, rump, throat, flank, upper breast, middle breast, and belly (Supplementary Figure 1). We measured each patch three times per individual and the spectrometer was calibrated using a white standard prior to measuring any new patch. We averaged the three measurements per patch per individual and removed electrical noise using functions implemented in the package “pavo” for R [38].

We quantified plumage coloration of six of the species belonging to the putative mimicry complex described by Prum (2014); we did not measure Gray-capped Flycatcher (*Myiozetetes granadensis*) or Yellow-throated Flycatcher (*Conopias parvus*) because their plumage patterns do not fulfill all of the “kiskadee-like” characteristics described above. We were unable to take measurements of White-ringed Flycatcher (*Conopias albovittatus*) because not enough specimens were available. We measured spectra from 10-11 specimens per species except for *P. inornata*, for which there where only seven specimens available, and *P. sulphuratus*, for which 19 specimens were measured. We used both female and male individuals and only measured undamaged specimens ≤ 50 years old [39]. We took measurements of 68 specimens (Supplementary Table 1), obtaining a total of 1,632 spectra.

#### Statistical and perceptual analysis

To determine whether species putatively involved in the mimicry complex are indeed indistinguishable from the perspectives of putative predators (raptors) or competitors (passerines), we addressed two questions following the approach described by Maia & White [40]: (1) Are the plumages of hypothetical models and mimics statistically distinct? and (2) Are their plumages perceptually different? We performed paired analysis between hypothetical models and mimics comparing coloration of each plumage patch using the averaged and noise-free spectra in the R package “pavo”[41] based on the receptor-noise model [42]. This model assumes thresholds for discrimination are imposed by receptor noise, which is dependent on the receptor type and its abundance in the retina [42,43]. The model allows one to estimate the distance between groups of points in a color space in units of “just noticeable differences” or JNDs [43]. If when comparing two colors the JND value is lower than 1, then those colors are predicted to be impossible to discriminate given the visual model and the chosen illuminant conditions [44–46].

To determine whether hypothetical models and mimics are statistically different in plumage coloration, we used permutation-based analyses of variance (PERMANOVAs) using perceptual color distances in the R package “vegan” [47]. We used 999 permutations and recorded the pseudo-f, the significance of the analysis (α=0.05), and the R^2^ [40]. To evaluate whether plumage patches showing statistical differences in reflectance are also perceptually distinguishable we did a bootstrap analysis to calculate a mean distance and a confidence interval in JNDs [40]. If two colors are statistically distinct and the lower bound of the bootstrapped confidence interval is higher than the established JND threshold value, then one can conclude that these colors are statistically distinct and perceptually different given a visual model [40]. Given that previous studies found that spectra separated by values ≤ 1 JND are impossible to distinguish and that even those with values ≤ 3 JNDs may be difficult to discriminate under bright light conditions [42,48], we chose an intermediate value of 2 JND as threshold to define two colors as perceptually distinguishable.

To assess statistical and perceptual differences from the perspective of raptors and tyrant flycatchers we performed PERMANOVAs and bootstraps assuming two alternative visual models. First, we used the “avg.v” model implemented in “pavo” which represents the standard violet-sensitive visual system; because there is no information available for Accipitriformes [49], we used receptor densities from the most closely related violet-sensitive relative, *Gallinula tenebrosa* (Rallidae) - *SWS1* **1**, *SWS2* **1.69**, *MWS* **2.10**, *LWS* **2.19**–[50]. We then used the “avg.uv” model representing the standard ultraviolet-sensitive visual system and used the default receptor densities *-SWS1* **1**, *SWS2* **2**, *MWS* **2**, *LWS* **4**- corresponding to *Leiothrix lutea* (Leothrichidae; [46]). We used a Weber fraction of 0.1 for both models [46] and the “bluesky” illuminant vector because our study species inhabit open areas [44,45].

We graphically examined plumage coloration using the Tetrahedral Color Space Model (TCS; [51,52]). The TCS model integrates data on sensitivity spectra of cones and luminance condition to transform reflectance spectra into points located in a tetrahedral color space, in which each corner represents the maximum stimulation for each cone type (*u*/*v*, *s*, *m*, *l*; [51,52]). Color spaces gave us an overview of plumage similarity between hypothetical model and mimic species given alternative visual discrimination models. We used the “vismodel”, “colspace” and “tetraplot” functions implemented in the R package “pavo” [41] and using the summary of the “colspace” result we recorded the total and relative color volume of each species as well as the *u*/*v*, *s*, *m* and *l* centroids to assess stimulation of each cone type. We also constructed reflectance curves (using the “aggplot” function) comparing each plumage patch of species involved in pairwise comparisons to visualize variation in hue (wavelength reflected). We corrected reflectance curves by mean brightness (“B2” measurement extracted from the “summary.rspec” result of the reflectance measurements of each patch; [41]) to visually assess differences based on hue and not brightness.

## RESULTS

### Can plumage similarity among flycatchers deceive putative competitors or predators?

As predicted by social mimicry hypotheses, we found some pairs of hypothetical model and mimic species, particularly those involving *M. pitangua*, to be indistinguishable from each other in plumage coloration (Table 1). Most hypothetical mimic species were perceptually indistinguishable from hypothetical model *M. pitangua* in the coloration of the eight patches we measured (JND values ≤2) in spite of some being statistically different from each other (Table 1 and Supplementary Table 2). *P. lictor* was distinguishable from *M. pitangua* in the flanks, but the discrimination value was very close to the discrimination threshold (2.06 JNDs and 2.10 JNDs for the UVS and VS models, respectively; Supplementary Table 2 and Supplementary Figure 3A). Moreover, all hypothetical mimics were indistinguishable from both hypothetical models in plumage from the crown, back, rump and throat (JND values≤2; Figure 2A, Figure 3A and Supplementary Table 3). Statistical and perceptual evaluation of the data were almost identical for the UVS and VS visual models (Table 1, Supplementary Table 2 and Supplementary Table 3), indicating that both predators and competitors might be deceived by the coloration of ventral plumage patches when considering *M. pitangua* as hypothetical model or by dorsal patches when considering either *M. pitangua* or *P. sulphuratus* as hypothetical models.

**Table 1.** Statistical and perceptual distinctiveness of plumage patches in parwise comparisons of species using the UVS and VS models. Patches that are statistically different are shown in yellow (α≥0.05 for the PERMANOVA). Patches that are perceptually different (JNDS>2) are shown with an asterisk.

| Patch/<br>Species | UVS |  |  |  |  |  |  |  | VS |  |  |  |  |  |  |  |
| --- | --- | --- | --- | --- | --- | --- | --- | --- | --- | --- | --- | --- | --- | --- | --- | --- |
|  | Upper<br>breast | Lower<br>breast | Belly | Flank | Throat | Crown | Back | Rump | Upper<br>breast | Lower<br>breast | Belly | Flank | Throat | Crown | Back | Rump |
| *M. pitangua*/<br>*M. cayanensis* |  |  |  |  |  |  |  |  |  |  |  |  |  |  |  |  |
| *M. pitangua*/<br>*M. similis* |  |  |  |  |  |  |  |  |  |  |  |  |  |  |  |  |
| *M. pitangua*/<br>*P. inornata* |  |  |  |  |  |  |  |  |  |  |  |  |  |  |  |  |
| *M. pitangua*/<br>*P. lictor* |  |  |  | * |  |  |  |  |  |  |  | * |  |  |  |  |
| *P. sulphuratus*/<br>*M. cayanensis* | * | * | * | * |  |  |  |  | * | * | * | * |  |  |  |  |
| *P. sulphuratus*/<br>*M. similis* | * | * | * | * |  |  |  |  | * | * | * | * |  |  |  |  |
| *P. sulphuratus*/<br>*P. inornata* | * | * | * | * |  |  |  |  | * | * | * | * |  |  |  |  |
| *P. sulphuratus*/<br>*P. lictor* | * | * | * | * |  |  |  |  | * | * | * | * |  |  |  |  |

**Figure 2.**
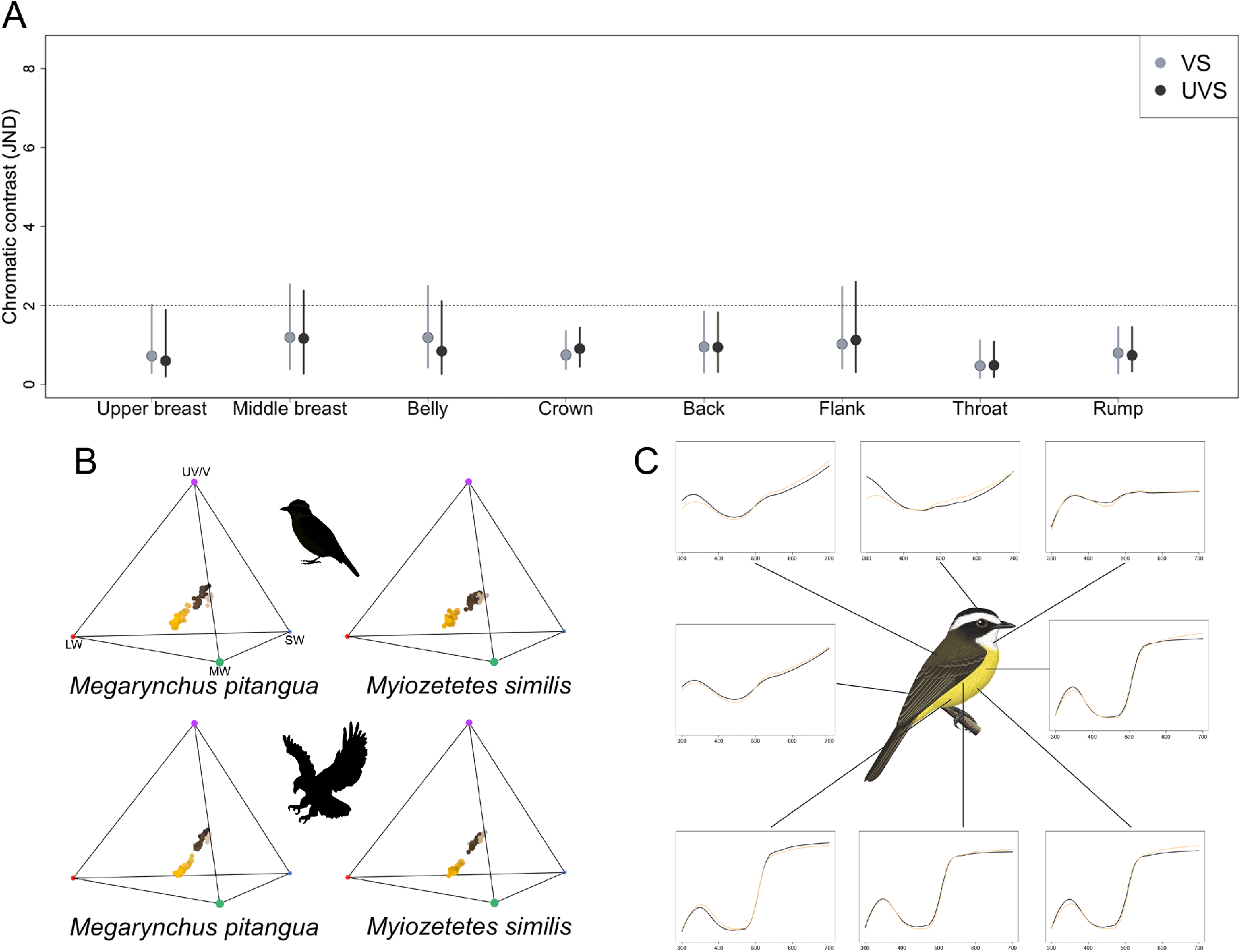
Example of a hypothetical pair of model (*Megarynchus pitangua*) and mimic (*Myiozetes similis*) species of flycatchers which we found are indistinguishable in plumage coloration under visual models describing discrimination abilities of putative competitors (UVS, passerines) and predators (VS, raptors). **A)** Color distances between species in units of chromatic contrast (just noticeable differences, JNDs) by plumage patch given the UVS (black) and the VS (gray) vision models. Points and bars are bootstrapped mean values and 95% confidence intervals, respectively. The dotted horizontal line indicates JND=2, below which colors are likely indistinguishable by birds. **B)** Coloration of plumage patches of each species in tetrahedral color space given UVS (top, i.e. competitors) and VS (bottom, i.e. predators) models; *uv/v, s, m* and *l* cone color channels are indicated in the first tetrahedron. Color spaces occupied by both species are highly similar given both vision models, but the color space volume varies between UVS and VS models. **C)** Reflectance curves for each plumage patch corrected by mean brilliance, with colors representing each of the two species being compared. There is little to no difference between model and mimic species reflectance curves in regards to hue. Illustration by Ayerbe-Quiñones [36] reproduced with permission from the author.

**Figure 3.**
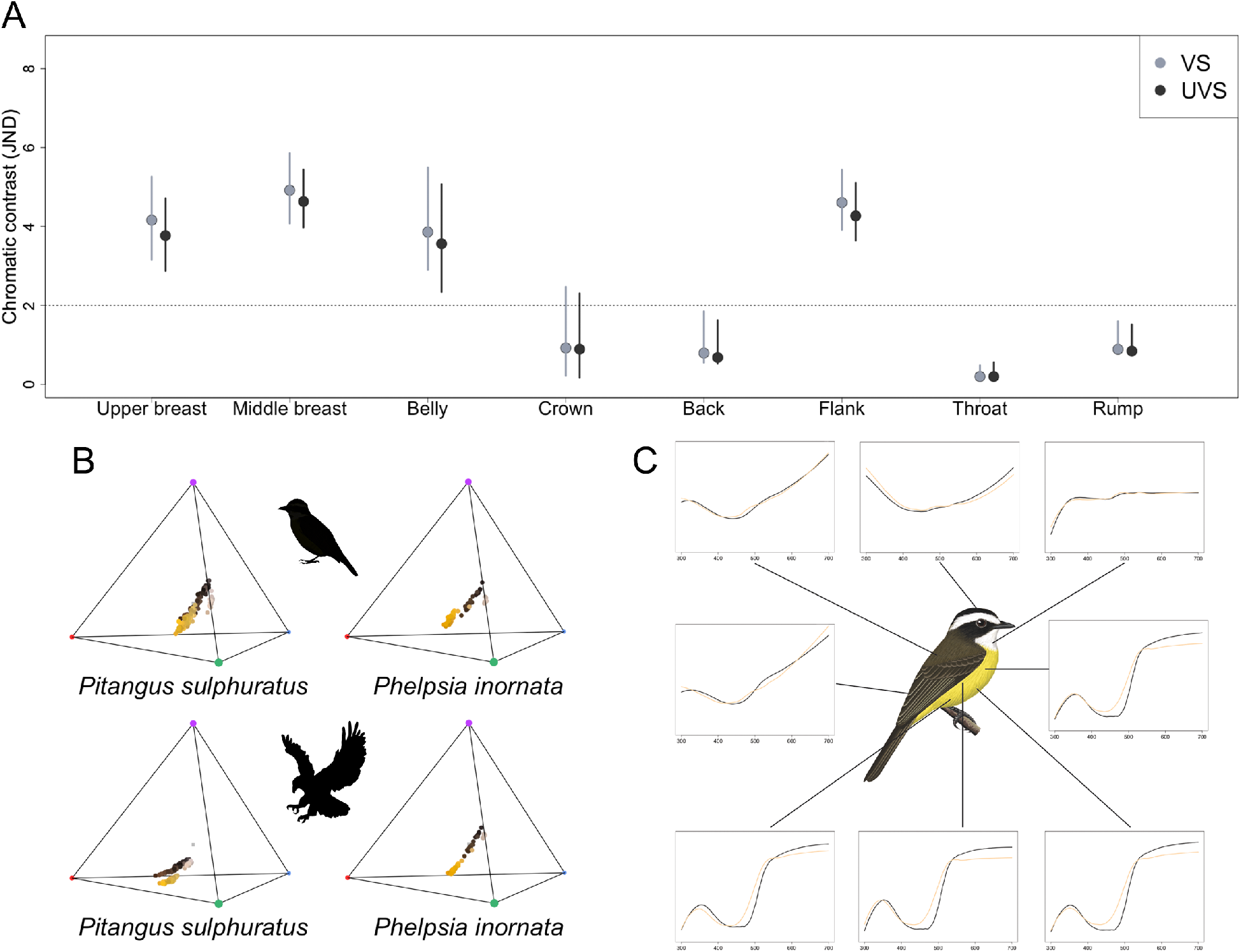
Example of a hypothetical pair of model (*Pitangus sulphuratus*) and mimic (*Phelpsia inornata’*) species of flycatchers which we found are distinguishable in plumage coloration under visual models of discrimination abilities of putative competitors (UVS, passerines) and predators (VS, raptors). **A)** Comparison of color distances (in units of chromatic contrast or just noticeable differences JNDs) by patch given the UVS (black) and the VS (gray) vision models. The dotted horizontal line indicates JND=2, below which the pair of colors is considered to be indistinguishable by birds. Points and bars indicate the bootstrapped mean value and 95% CI’s respectively. **B)** Distribution of the color volume of each species in the tetrahedral color space using UVS and VS models. Volumes occupied by individual species differ between vision models; for a given vision model, color spaces occupied by each species are distinct. **C)** Reflectance curves for each plumage patch corrected by mean brilliance, with curve colors representing each species being compared. There is a marked difference between model and mimic species ventral reflectance curves in regards to hue around 500nm. Illustrations by Ayerbe-Quiñones [36] reproduced with permission from the author.

Resemblance between *M. pitangua* and hypothetical mimics exists because although there are differences in brilliance of all plumage patches, the hue reflected by each patch is highly similar between species (Figure 2C). Although descriptive variables of the plumage (i.e usml centroids and total and relative volumes) of each species vary between the two visual models (Supplementary Table 4), this variation is not relevant when evaluating discriminability between hypothetical model and mimic species. As a graphical example, distribution of the points in the tetrahedral representation were very similar between the two visual models, except that measurements from a given individual occupied a larger volume in the UVS model (Figure 2B).

Conversely, we found that pairs of hypothetical model and mimic species involving *P. sulphuratus* are distinguishable in plumage, particularly on ventral patches (Table 1). All hypothetical mimics were perceptually distinguishable from hypothetical model *P. sulphuratus* in plumage of the upper breast, middle breast, belly and flank patches (JND values > 2; Table 1 and Figure 3A). Underpart patches were statistically and perceptually different in all comparisons (Table 1). Color dissimilarity between *P. sulphuratus* and hypothetical mimics is illustrated by difference in the wavelengths reflected around 500nm in underpart patches (Figure 3C) and by differences in the stimulation values of the *s* color cone (Supplementary Table 5). Statistical and perceptual evaluation of the data were almost identical both for the UVS and VS models, indicating that hypothetical mimic species are distinguishable from *P. sulphuratus* by model species, smaller passerine species, and predatory raptors. Similar results are evident in tetrahedral representations, except that, again, in the UVS model measurements from a given individual occupied a larger volume (Figure 3B).

## DISCUSSION

Hypotheses posed to account for phenotypic convergence involving mimicry have seldom been assessed while considering the visual systems of receivers. For example, recent work on *Heliconius* butterflies involved in Müllerian mimicry rings revealed that coloration patterns of comimics appearing similar to the human eye are actually distinct given the visual abilities of butterflies, yet may be indistinguishable by some of their avian predators, particularly those with VS visual systems [53]. That predators indeed perceive putative comimics as similar -and associate their appearance with unpalatability-validates a core assumption of the hypothesis that shared color patterns confer adaptive benefits which result in convergence in aposematic coloration among chemically defended butterfly species [54]. Likewise, that putative mimic and model species are indistinguishable by competitors or predators is critical for a number of hypotheses posed to account for plumage convergence involving mimicry in birds to be plausible [7,13,16–22]. Prior to our study, however, this critical assumption had seldom been critically examined.

Although convergence in plumage patterns is widespread across birds [7,8,10–13,16–19,21,55], few studies have assessed the mechanisms underlying this phenomenon. For example, convergence has been documented in birds which may engage in mimicry including toucans [56], friarbirds and orioles [8], and woodpeckers [13,15]. However, the extent to which alternative hypotheses involving mimicry may account for convergence in these groups is largely unknown. We here assessed the plausibility of mimicry hypotheses using spectrophotometric data in six distantly related, but strikingly similar tyrant-flycatcher species. Although we found some evidence consistent with mimicry hypotheses, some of our results indicate that at least part of the explanations for the striking phenotypic similarity due to convergence among species of flycatchers may require reconsideration.

Our results revealed that all hypothetical mimic species are indistinguishable from hypothetical model species in coloration of dorsal plumage patches given model of visual discrimination resembling that of raptors (VS model). This result supports the hypothesis that mimicry in birds may arise as an antipredator strategy [16], which predicts that plumages should be indistinguishable to predators given their visual system. Moreover, mimicry should be more precise in plumage patches used by predators as cues to select prey [16]. The main predators of adult songbirds, including tyrant flycatchers, are likely diurnal raptors [27,28,57,58], which often observe prey from long distances while perched on treetops [59] and may choose odd individuals relative to their background [24,32]. Consequently, similarity in dorsal coloration in species that forage together or use different strata of the same trees may arguably create a sense of homogeneity and thereby be adaptive to avoid attacks from predators approaching from above.

Wallace [60,61] and later Diamond [20] were amazed by the striking similarity in plumage between Australian orioles (genus *Oriolus*, family Oriolidae) and friarbirds (genus *Philemon*, family Meliphagidae). Wallace first claimed such similarity was a case of visual mimicry, but no study on the subject was done until Diamond [20] posited that visual mimicry may serve to escape attack from larger model species or to deceive smaller species and scare them off only by appearance. Prum & Samuelson [19] and Prum [7] further expanded on the first idea by positing the ISDM hypothesis and outlining its predictions. A recent analysis assessing the ISDM on orioles and friarbirds using phylogenetic methods suggested that orioles indeed appear to mimic larger-bodied friarbirds [8], but there is no information about the species being deceived in this system. In principle, ISDM may also apply to kiskadee-like flycatchers because existing body size data supports the prediction that hypothetical model species are larger in body mass (i.e. at least 30g heavier) than hypothetical mimic species [34]. The additional prediction that models are socially dominant over mimics has not been tested quantitatively, but several observations exist of both hypothetical model species scaring off hypothetical mimics from foraging grounds (David Ocampo, Santiago Rosado, and Oscar Laverde, pers. comm.).

A critical additional prediction of the ISDM hypothesis is that visual deception based on convergent coloration should be physiologically plausible at ecologically relevant visual distances among individuals [7]. We found partial support for this prediction. On one hand, our results show that hypothetical mimics were perceptually distinguishable from hypothetical model *P. sulphuratus* in the coloration of the upper breast, middle breast, abdomen and flank patches using the UVS model. Considering that underpart patches are visually relevant when two species engage physically in interference competition [62], our analyses reject the proposition that visual deception is physiologically possible when assuming *P. sulphuratus* as hypothetical model. This result is consistent with previous work in other birds with striking similarity to the human eye: putatively mimetic Downy Woodpeckers (*Picoides pubescens*) do not experience reduced aggression from hypothetical model Hairy Woodpeckers (*Picoides villosus*), implying lack of deception [13]. Because Downy Woodpeckers are more dominant over other bird species than expected based on their body size, convergence in plumage with Hairy Woodpeckers may instead have evolved to deceive smaller third-party species [13,20], a hypothesis yet to be tested in kiskadee-like flycathers resembling *P. sulphuratus*. On the other hand, we found that most hypothetical mimics are perceptually indistinguishable from *M. pitangua* in ventral plumage patches. Perceptual similarity under the UVS model indicates that *M. pitangua* might be deceived by hypothetical mimics, misidentify them as conspecifics, and thus split resources with them owing to reduced aggression. Alternatively, other passerines might also be deceived by hypothetical mimics, misidentify them as *M. pitangua* individuals, and therefore withdraw from an aggressive interaction. Consequently, our results are consistent with mimicry hypotheses that imply deception of either putative models or smaller passerine competitors [7,19–21] when considering *M. pitangua* as the putative model. We are unable to fully discriminate between the two hypotheses with our results, but we agree with Leighton et al. [13] that visual deception of hypothetical models seems unlikely because individuals are expected to be adept at identifying conspecifics given its importance for competition and successful breeding. In addition, plumage is likely not the only cue that birds employ to recognize conspecifics in the field and *M. pitangua* is structurally different from its potential mimics owing to its massive bill, which is readily recognizable even by human observers. Alternatively, because selective pressures to identify individuals which are not predators, prey or strong competitors are likely reduced, visual deception of species that are neither hypothetical mimics or models may be more likely [13,20].

Ours is the first study to assess the plausibility of mimicry hypotheses in birds using spectrophotometric measurements of plumage, and evaluating the data with statistical and perceptual analysis [40] given two avian visual models. Additional work is required to further evaluate hypotheses accounting for plumage convergence. For instance, although our study species overlap in geographic range, diet and foraging strategies [37,63,64], very little is known about interactions among them, and the extent to which hypothetical models are indeed deceived by hypothetical mimics should be evaluated through behavioral observations and experiments. Likewise, field studies are required to assess whether predators such as raptors are indeed deceived by putative models and mimics to escape predation. In addition, there is no knowledge of how perception of color may vary with distance between individuals or of how to account for distances over which individuals interact in the field when analyzing spectrophotometric data. Hence, we do not know precisely how likely deception is at ecologically relevant distances, an important condition for ISDM [7]. For example, while some hypothetical models may be distinguishable by hypothetical mimics upon inspection at close distances, hypothetical mimic species may still be able to deceive hypothetical models from greater distances [13].

A caveat of our analyses is that we modelled the visual system of predators based only on the species phylogenetically closest to accipitrid raptors for which information was available, namely *Gallinula tenebrosa*. However, *G. tenebrosa* is a non-raptorial bird and we lack information on how closely photoreceptor densities and peak sensitivites resemble that of acciptrids. Other raptors with likely different vision systems (i.e. falcons, Falconiformes) also prey upon flycatchers, but specific models for such predators are also lacking. Likewise, we modelled the visual system of our study species of flycatchers given existing standard models of passerine UVS vision based on other species. More definitive tests of the hypothesis that similarities among species of flycatchers with kiskadee-like plumage may deceive their avian predators or putative competitors thus await additional consideration involving the use of visual models developed specifically for predatory species and for the birds we studied. This is because variation among and within species in visual abilities may exist [65,66] and because differences in unstudied traits such as photoreceptor densities may have large consequences on the ability of species to discriminate between similar colors [49]. Given these caveats and that information on specific visual models of the species involved in our study are measing we opted to set a conservative threshold (2 JNDs) to define two colors as perceptually different.

In conclusion, perceptual similarity of the crown, back and rump patches among species is consistent with the hypothesis that predation by visually oriented predators approaching their prey from above may have favored convergence in plumage in kiskadee-like tyrant flycatchers [16]. Perceptual similarity in ventral patches suggests that deception involved in competitive interactions with *M. pitangua*, but not with *P. sulphuratus*, may also have favored convergence [7,19–21]. Future studies should focus on gathering behavioral data to characterize competitive and predator-prey interactions among species potentially involved in social mimicry. Assessing how other factors like climate, habitat and development shape the evolution of plumage would allow for a comprehensive understanding of the mechanisms underlying convergence.

## Acknowledgments

We thank Museo de Historia Natural de la Universidad de los Andes (ANDES), Instituto de Ciencias Naturales de la Universidad Nacional (ICN), and Instituto de Investigación de Recursos Biológicos Alexander von Humboldt (IAvH) for allowing us take spectrophotometric measurements of museum specimens. We are specially grateful to David Slager for helping structure the original idea, guidance in the first development phase of this work, and for thoughful comments on the manuscript. We thank Gustavo Bravo, Michael Harvey, and colleagues for sharing information on their flycatcher phylogeny. Members of the Laboratorio de Biología Evolutiva de Vertebrados provided insightful comments and support during the development of this project. Special thanks to Natasha Bloch for comments on the manuscript, help with the reflectance curves and nutritive conversations about the project. Finally, thanks to Laura Cespedes-Arias for providing guidance throughout the process of data collection and analysis.

---

**Supplementary Table 1.**
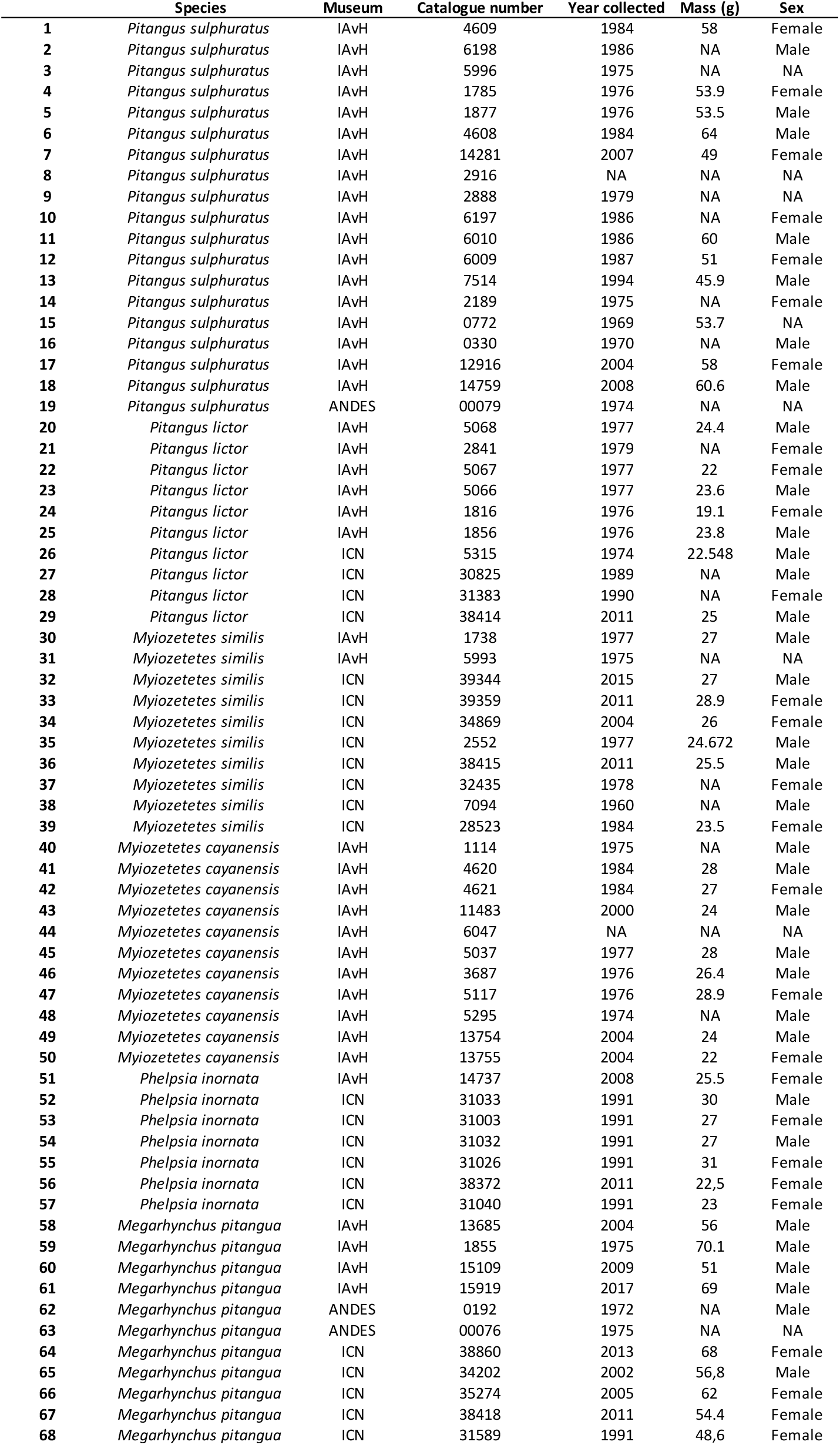
Complete specimen information

**Supplementary Table 2.** Pseudo-f, R^2^ and significance (α=0.05) for the PERMANOVA using the UVS and VS models. Patches that are statistically different are bolded and highlighted in gray.

| Patch/<br>Species | UVS |  |  |  |  |  |  |  | VS |  |  |  |  |  |  |  |
| --- | --- | --- | --- | --- | --- | --- | --- | --- | --- | --- | --- | --- | --- | --- | --- | --- |
|  | Upper<br>breast | Lower<br>breast | Belly | Flank | Throat | Crown | Back | Rump | Upper<br>breast | Lower<br>breast | Belly | Flank | Throat | Crown | Back | Rump |
| *M. pitangua/*<br>*M. cayanensis* | 0.484<br>0.025<br>(0.656) | 1.818<br>0.092<br>(0.164) | 1.249<br>0.072<br>(0.305) | 1.220<br>0.075<br>(0.297) | 0.699<br>0.037<br>(0.558) | **5.974**<br>**0.249**<br>**(0.013)** | **6.190**<br>**0.256**<br>**(0.012)** | **13.911**<br>**0.436**<br>**(0.001)** | 0.376<br>0.019<br>(0.706) | 1.915<br>0.096<br>(0.154) | 1.285<br>0.074<br>(0.275) | 1.560<br>0.094<br>(0.202) | 0.134<br>0.007<br>(0.912) | **5.416**<br>**0.231**<br>**(0.021)** | **6.485**<br>**0.265**<br>**(0.009)** | **16.260**<br>**0.475**<br>**(0.001)** |
| *M. pitangua/*<br>*M. similis* | 1.094<br>0.054<br>(0.322) | 1.601<br>0.082<br>(0.197) | 2.133<br>0.111<br>(0.133) | 1.255<br>0.062<br>(0.266) | 1.825<br>0.102<br>(0.196) | **4.277**<br>**0.192**<br>**(0.014)** | 2.354<br>0.122<br>(0.120) | 1.568<br>0.080<br>(0.232) | 1.376<br>0.068<br>(0.255) | 1.491<br>0.076<br>(0.236) | 2.665<br>0.136<br>(0.095) | 1.021<br>0.051<br>(0.342) | 1.945<br>0.108<br>(0.167) | **3.471**<br>**0.162**<br>**(0.033)** | 2.584<br>0.132<br>(0.089) | 2.575<br>0.125<br>(0.088) |
| *M. pitangua/*<br>*P. inornata* | 0.485<br>0.029<br>(0.617) | **5.160**<br>**0.256**<br>**(0.016)** | 1.685<br>0.107<br>(0.135) | 2.322<br>0.127<br>(0.115) | 1.621<br>0.098<br>(0.181) | 0.702<br>0.045<br>(0.662) | 1.850<br>0.110<br>(0.174) | **5.604**<br>**0.259**<br>**(0.001)** | 0.322<br>0.020<br>(0.709) | **5.520**<br>**0.269**<br>**0.011** | 2.095<br>0.130<br>(0.062) | 2.368<br>0.129<br>(0.104) | 1.410<br>0.086<br>(0.237) | 0.659<br>0.042<br>(0.689) | 2.041<br>0.120<br>(0.151) | **6.949**<br>**0.303**<br>**(0.001)** |
| *M. pitangua/*<br>*P. lictor* | 0.940<br>0.047<br>(0.388) | **12.910**<br>**0.418**<br>**(0.001)** | 1.341<br>0.069<br>(0.255) | **12.497**<br>**0.397**<br>**(0.001)** | 2.242<br>0.117<br>(0.116) | 0.703<br>0.038<br>(0.499) | **7.729**<br>**0.300**<br>**(0.003)** | **6.176**<br>**0.245**<br>**(0.003)** | 0.970<br>0.049<br>(0.366) | **12.922**<br>**0.418**<br>**(0.002)** | **3.702**<br>**0.171**<br>**(0.029)** | **12.528**<br>**0.397**<br>**(0.003)** | 1.543<br>0.083<br>(0.220) | 0.703<br>0.038<br>(0.538) | **7.430**<br>**0.292**<br>**(0.001)** | **6.671**<br>**0.260**<br>**(0.001)** |
| *P. sulphuratus/*<br>*M. cayanensis* | **24.069**<br>**0.471**<br>**(0.001)** | **18.423**<br>**0.415**<br>**(0.001)** | **24.886**<br>**0.531**<br>**(0.001)** | **16.045**<br>**0.422**<br>**(0.001)** | 1.173<br>0.043<br>(0.320) | **12.877**<br>**0.331**<br>**(0.002)** | 1.192<br>0.044<br>(0.292) | 1.238<br>0.047<br>(0.302) | **20.826**<br>**0.435**<br>**(0.001)** | **17.869**<br>**0.407**<br>**(0.001)** | **22.546**<br>**0.506**<br>**(0.001)** | **16.298**<br>**0.426**<br>**(0.002)** | 1.592<br>0.058<br>(0.216) | **11.364**<br>**0.304**<br>**(0.003)** | 1.574<br>0.057<br>(0.197) | 1.481<br>0.056<br>(0.217) |
| *P. sulphuratus/*<br>*M. similis* | **47.573**<br>**0.638**<br>**(0.001)** | **34.200**<br>**0.568**<br>**(0.001)** | **19.795**<br>**0.463**<br>**(0.001)** | **28.591**<br>**0.524**<br>**(0.001)** | **6.320**<br>**0.208**<br>**(0.003)** | **9.642**<br>**0.271**<br>**(0.004)** | 1.108<br>0.042<br>(0.303) | **4.239**<br>**0.145**<br>**(0.024)** | **50.912**<br>**0.653**<br>**(0.001)** | **38.595**<br>**0.597**<br>**(0.001)** | **22.902**<br>**0.499**<br>**(0.001)** | **34.733**<br>**0.572**<br>**(0.001)** | **6.887**<br>**0.223**<br>**(0.005)** | **8.207**<br>**0.240**<br>**(0.005)** | 1.234<br>0.047<br>(0.270) | **4.19**<br>**0.144**<br>**(0.013)** |
| *P. sulphuratus/*<br>*P. inornata* | **25.581**<br>**0.516**<br>**(0.001)** | **47.832**<br>**0.675**<br>**(0.001)** | **16.569**<br>**0.453**<br>**(0.001)** | **33.133**<br>**0.590**<br>**(0.001)** | 0.683<br>0.029<br>(0.529) | 1.151<br>0.048<br>(0.332) | 0.179<br>0.008<br>(0.882) | 1.003<br>0.042<br>(0.363) | **28.973**<br>**0.547**<br>**(0.001)** | **48.110**<br>**0.677**<br>**(0.001)** | **18.993**<br>**0.487**<br>**(0.001)** | **41.564**<br>**0.644**<br>**(0.001)** | 0.865<br>0.036<br>(0.423) | 1.037<br>0.043<br>(0.363) | 0.280<br>0.012<br>(0.803) | 1.106<br>0.046<br>(0.345) |
| *P. sulphuratus/*<br>*P. lictor* | **32.470**<br>**0.546**<br>**(0.001)** | **89.794**<br>**0.775**<br>**(0.001)** | **54.405**<br>**0.694**<br>**(0.001)** | **93.284**<br>**0.782**<br>**(0.001)** | 0.338<br>0.013<br>(0.723) | 0.702<br>0.026<br>(0.461) | 1.620<br>0.059<br>(0.188) | 1.219<br>0.045<br>(0.294) | **33.929**<br>**0.557**<br>**(0.001)** | **89.464**<br>**0.775**<br>**(0.001)** | **64.579**<br>**0.729**<br>**(0.001)** | **102.971**<br>**0.798**<br>**(0.001)** | 0.152<br>0.006<br>(0.897) | 0.541<br>0.020<br>(0.543) | 2.016<br>0.072<br>(0.138) | 1.094<br>0.040<br>(0.357) |

**Supplementary Table 3.** Upper, mean and lower values of JND resulting of the bootstrap analysis using the UVS and VS models. Patches that are perceptually different are bolded and highlighted in gray.

| Patch/<br>Species | UVS |  |  |  |  |  |  |  | VS |  |  |  |  |  |  |  |
| --- | --- | --- | --- | --- | --- | --- | --- | --- | --- | --- | --- | --- | --- | --- | --- | --- |
|  | Upper<br>breast | Lower<br>breast | Belly | Flank | Throat | Crown | Back | Rump | Upper<br>breast | Lower<br>breast | Belly | Flank | Throat | Crown | Back | Rump |
| *M. pitangua*/<br>*M. cayanensis* | 1.982 | 2.761 | 1.561 | 3.721 | 0.684 | 2.101 | 2.674 | 2.822 | 2.035 | 3.054 | 1.700 | 4.382 | 0.626 | 1.983 | 2.671 | 2.867 |
|  | 0.689 | 1.355 | 0.503 | 1.487 | 0.185 | 1.293 | 1.726 | 2.034 | 0.736 | 1.492 | 0.698 | 1.863 | 0.080 | 1.235 | 1.724 | 2.094 |
|  | 0.285 | 0.353 | 0.160 | 0.208 | 0.075 | 0.462 | 0.762 | 1.272 | 0.420 | 0.471 | 0.227 | 0.374 | 0.053 | 0.448 | 0.731 | 1.316 |
| *M. pitangua*/<br>*M. similis* | 1.890 | 2.374 | 2.110 | 2.603 | 1.080 | 1.441 | 1.831 | 1.459 | 2.018 | 2.532 | 2.495 | 2.477 | 1.113 | 1.357 | 1.847 | 1.459 |
|  | 0.595 | 1.164 | 0.841 | 1.124 | 0.484 | 0.905 | 0.943 | 0.739 | 0.723 | 1.192 | 1.187 | 1.021 | 0.471 | 0.747 | 0.948 | 0.794 |
|  | 0.201 | 0.269 | 0.260 | 0.308 | 0.190 | 0.453 | 0.311 | 0.329 | 0.288 | 0.396 | 0.423 | 0.405 | 0.161 | 0.390 | 0.305 | 0.276 |
| *M. pitangua*/<br>*P. inornata* | 2.003 | 3.298 | 3.795 | 2.635 | 0.731 | 1.923 | 2.226 | 2.758 | 1.984 | 3.425 | 4.484 | 2.751 | 0.787 | 1.918 | 2.140 | 2.678 |
|  | 0.719 | 2.078 | 1.151 | 1.381 | 0.470 | 0.421 | 1.005 | 1.730 | 0.492 | 2.159 | 1.697 | 1.433 | 0.370 | 0.437 | 0.998 | 1.796 |
|  | 0.330 | 1.044 | 0.461 | 0.484 | 0.330 | 0.118 | 0.220 | 0.951 | 0.354 | 0.979 | 0.893 | 0.838 | 0.155 | 0.118 | 0.226 | 1.064 |
| *M. pitangua*/<br>*P. lictor* | 3.054 | 4.207 | 1.432 | **4.658** | 1.175 | 1.300 | 2.644 | 2.655 | 3.149 | 4.401 | 1.842 | **4.882** | 1.005 | 1.423 | 2.593 | 2.736 |
|  | 1.145 | 3.028 | 0.757 | **3.381** | 0.717 | 0.189 | 1.750 | 1.731 | 1.376 | 3.129 | 1.310 | **3.525** | 0.534 | 0.159 | 1.694 | 1.781 |
|  | 0.309 | 1.918 | 0.401 | **2.064** | 0.568 | 0.084 | 0.922 | 0.911 | 0.831 | 1.971 | 0.943 | **2.104** | 0.375 | 0.101 | 0.815 | 0.947 |
| *P. sulphuratus*/<br>*M. cayanensis* | **5.102** | **4.783** | **4.889** | **5.045** | 1.532 | 1.918 | 1.609 | 2.364 | **5.464** | **5.058** | **5.086** | **5.354** | 1.595 | 1.793 | 1.777 | 2.222 |
|  | **4.308** | **3.902** | **3.752** | **3.996** | 1.088 | 1.353 | 0.978 | 1.518 | **4.566** | **4.183** | **4.108** | **4.364** | 0.993 | 1.173 | 1.060 | 1.441 |
|  | **3.446** | **3.069** | **2.804** | **3.041** | 0.772 | 0.755 | 0.780 | 0.970 | **3.686** | **3.545** | **3.440** | **3.531** | 0.541 | 0.577 | 0.833 | 1.060 |
| *P. sulphuratus*/<br>*M. similis* | **4.257** | **4.787** | **4.916** | **6.036** | 0.989 | 2.568 | 2.258 | 1.520 | **4.495** | **5.023** | **5.310** | **6.572** | 1.037 | 2.528 | 2.327 | 1.639 |
|  | **3.282** | **3.516** | **3.985** | **4.148** | 0.568 | 1.793 | 1.197 | 0.726 | **3.404** | **3.703** | **4.190** | **4.473** | 0.552 | 1.739 | 1.336 | 0.801 |
|  | **2.346** | **2.605** | **3.094** | **2.292** | 0.297 | 0.947 | 0.567 | 0.531 | **2.369** | **2.604** | **3.091** | **2.616** | 0.268 | 0.930 | 0.586 | 0.580 |
| *P. sulphuratus*/<br>*P. inornata* | **4.817** | **5.617** | **5.177** | **5.077** | 0.573 | 2.464 | 1.670 | 1.542 | **5.224** | **5.893** | **5.606** | **5.518** | 0.502 | 2.482 | 1.898 | 1.670 |
|  | **3.782** | **4.709** | **3.444** | **4.244** | 0.160 | 0.921 | 0.686 | 0.837 | **4.227** | **5.021** | **4.129** | **4.768** | 0.219 | 0.949 | 0.817 | 0.953 |
|  | **2.801** | **3.959** | **2.024** | **3.426** | 0.082 | 0.180 | 0.516 | 0.760 | **3.372** | **4.275** | **3.193** | **4.145** | 0.154 | 0.240 | 0.600 | 0.867 |
| *P. sulphuratus*/<br>*P. lictor* | **6.431** | **6.549** | **5.454** | **7.150** | 0.877 | 1.769 | 2.292 | 1.581 | **6.843** | **6.962** | **5.913** | **7.739** | 0.683 | 1.887 | 2.350 | 1.695 |
|  | **4.893** | **5.724** | **4.615** | **6.190** | 0.116 | 0.686 | 1.221 | 0.869 | **5.203** | **6.012** | **5.194** | **6.740** | 0.015 | 0.670 | 1.390 | 0.963 |
|  | **3.352** | **4.784** | **3.799** | **5.200** | 0.060 | 0.163 | 0.564 | 0.761 | **3.827** | **5.158** | **4.581** | **5.810** | 0.042 | 0.125 | 0.695 | 0.869 |

**Supplementary Figure 1.**
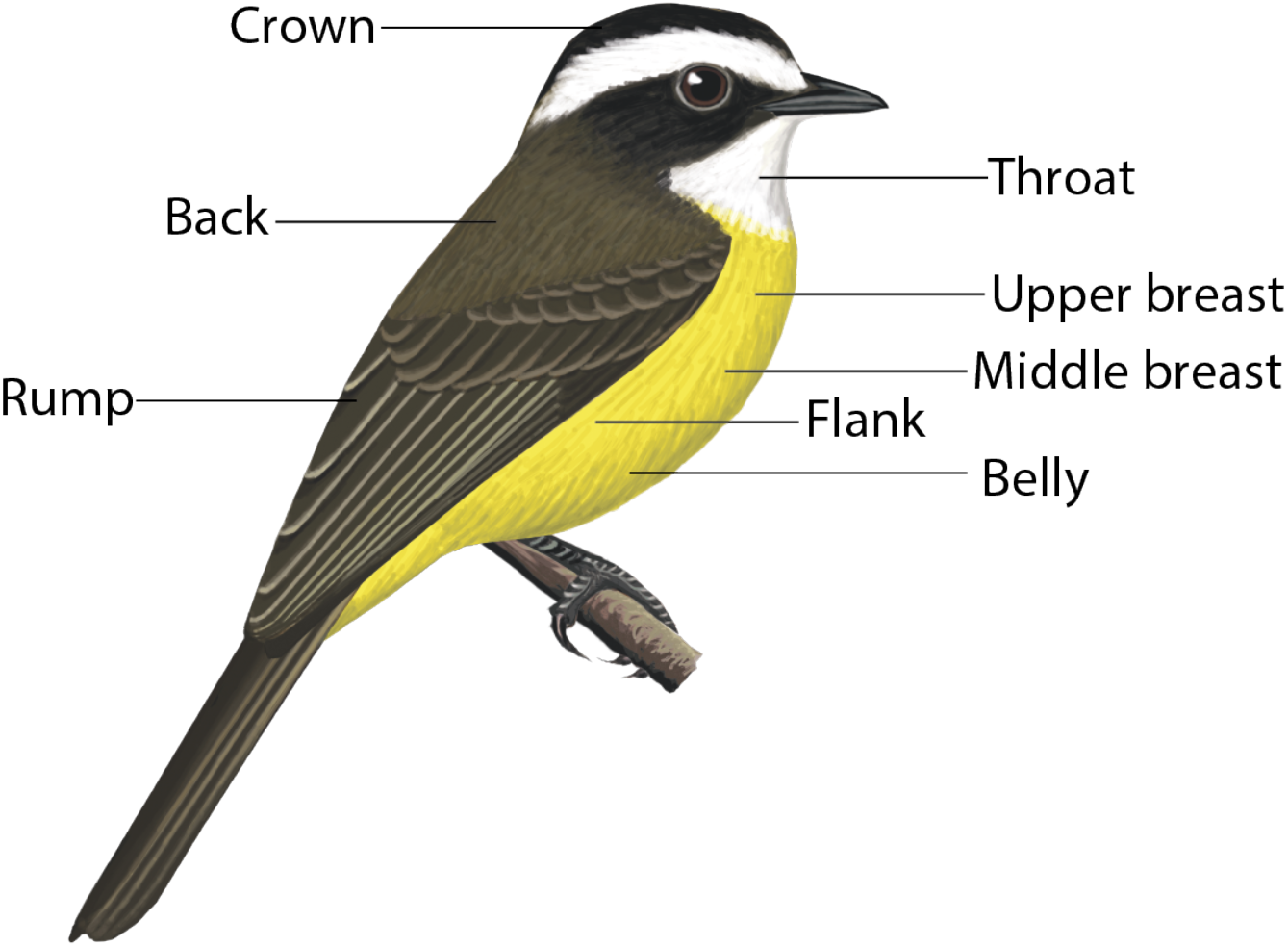
Plumage patches measured to characterize coloration and compare plumage among species of “kiskadee-like” flycatchers. Illustration by Ayerbe-Quiñones [36] reproduced with permission from the author.

**Supplementary Figure 2.**
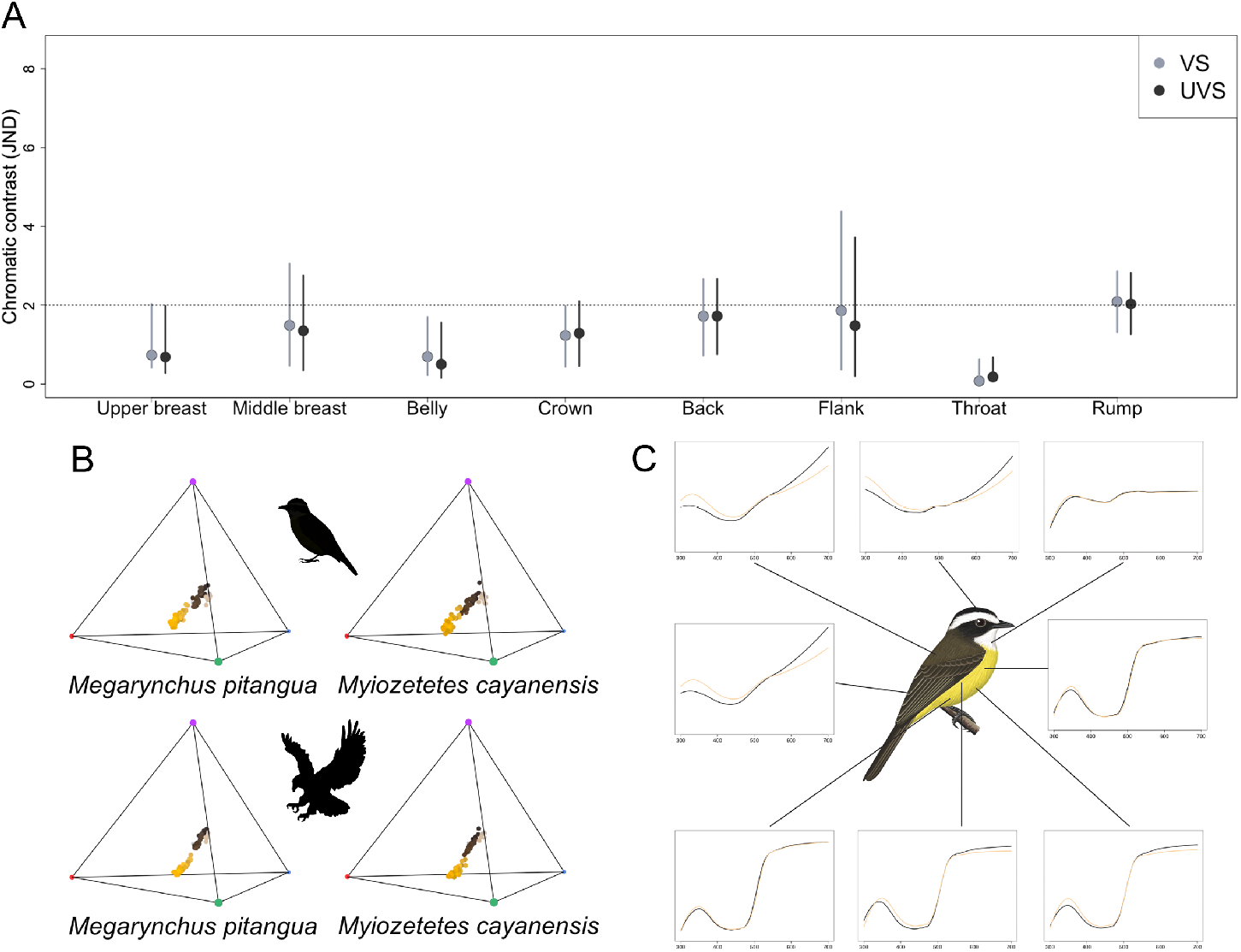
A hypothetical pair of model (*Megarynchus pitangua*) and mimic (*Myiozetetes cayanensis*) species of flycatchers which we found are indistinguishable in plumage coloration under visual models describing discrimination abilities of putative competitors (UVS, passerines) and predators (VS, raptors). A) Color distances between species in units of chromatic contrast (just noticeable differences, JNDs) by plumage patch given the UVS (black) and the VS (gray) vision models. Points and bars are bootstrapped mean values and 95% confidence intervals, respectively. The dotted horizontal line indicates JND=2, below which colors are likely indistinguishable by birds. B) Coloration of plumage patches of each species in tetrahedral color space given UVS (top, i.e. competitors) and VS (bottom, i.e. predators) models. Color spaces occupied by both species are highly similar given both vision models, but the color space volume varies between UVS and VS models. C) Reflectance curves for each plumage patch corrected by mean brilliance, with curve colors representing each of the two species being compared. There is little to no difference between model and mimic species reflectance curves in regards to hue. Illustration by Ayerbe-Quiñones [36] reproduced with permission from the author.

**Supplementary Figure 3.**
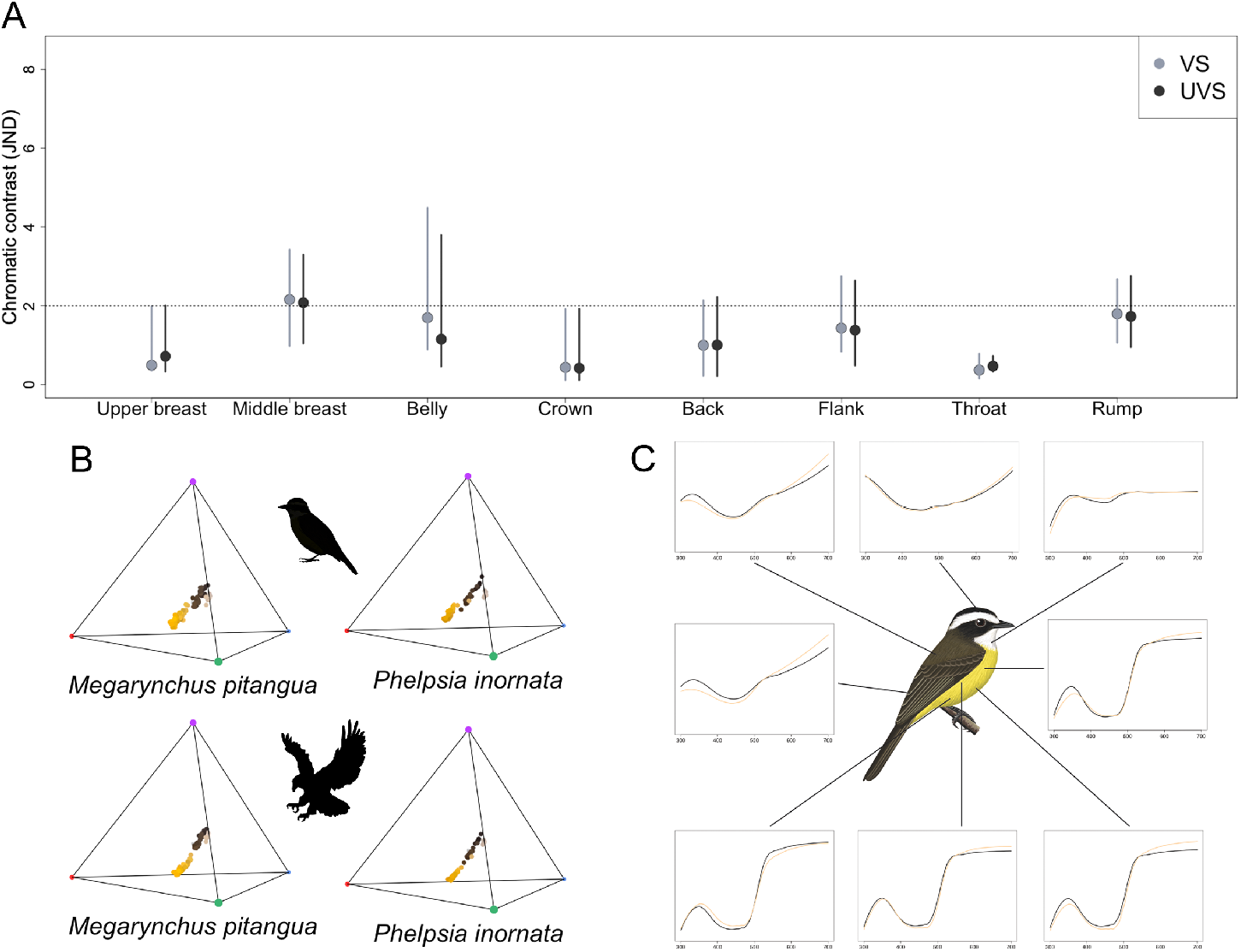
A hypothetical pair of model (*Megarynchus pitangua*) and mimic (*Phelpsia inornata*) species of flycatchers which we found are indistinguishable in plumage coloration under visual models describing discrimination abilities of putative competitors (UVS, passerines) and predators (VS, raptors). **A)** Color distances between species in units of chromatic contrast (just noticeable differences, JNDs) by plumage patch given the UVS (black) and the VS (gray) vision models. Points and bars are bootstrapped mean values and 95% confidence intervals, respectively. The dotted horizontal line indicates JND=2, below which colors are likely indistinguishable by birds. **B)** Coloration of plumage patches of each species in tetrahedral color space given UVS (top, i.e. competitors) and VS (bottom, i.e. predators) models. Color spaces occupied by both species are highly similar given both vision models, but the color space volume varies between UVS and VS models. **C)** Reflectance curves for each plumage patch corrected by mean brilliance, with curve colors representing each of the two species being compared. There is little to no difference between model and mimic species reflectance curves in regards to hue. Illustration by Ayerbe-Quiñones [36] reproduced with permission from the author.

**Supplementary Figure 4.**
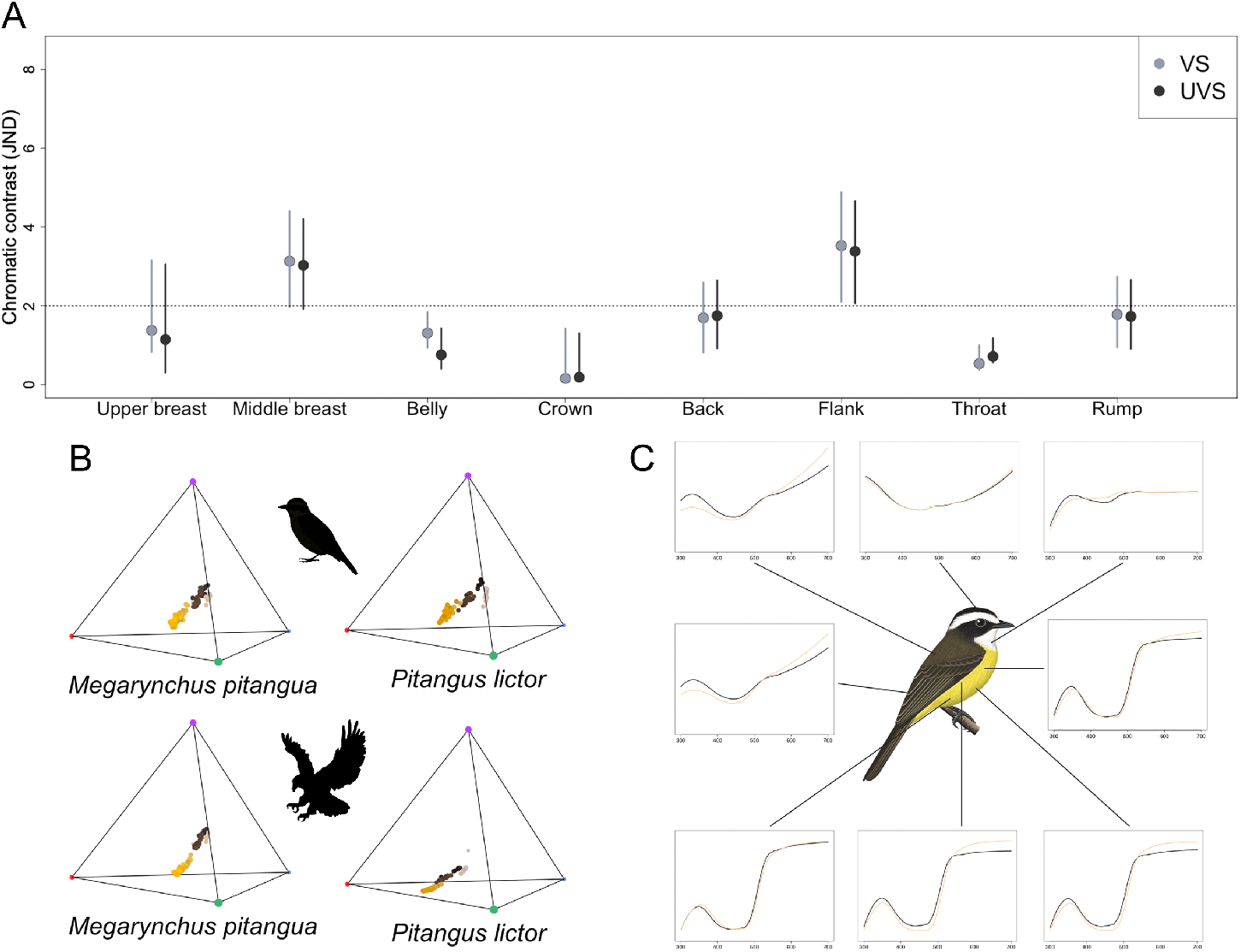
A hypothetical pair of model (*Megarynchus pitangua*) and mimic (*Pitangus lictor*) species of flycatchers which we found are indistinguishable in plumage coloration under visual models describing discrimination abilities of putative competitors (UVS, passerines) and predators (VS, raptors). **A)** Color distances between species in units of chromatic contrast (just noticeable differences, JNDs) by plumage patch given the UVS (black) and the VS (gray) vision models. Points and bars are bootstrapped mean values and 95% confidence intervals, respectively. The dotted horizontal line indicates JND=2, below which colors are likely indistinguishable by birds. **B)** Coloration of plumage patches of each species in tetrahedral color space given UVS (top, i.e. competitors) and VS (bottom, i.e. predators) models. Color spaces occupied by both species are highly similar given both vision models, but the color space volume varies between UVS and VS models. **C)** Reflectance curves for each plumage patch corrected by mean brilliance, with curve colors representing each of the two species being compared. There is little to no difference between model and mimic species reflectance curves in regards to hue. Illustration by Ayerbe-Quiñones [36] reproduced with permission from the author.

**Supplementary Figure 5.**
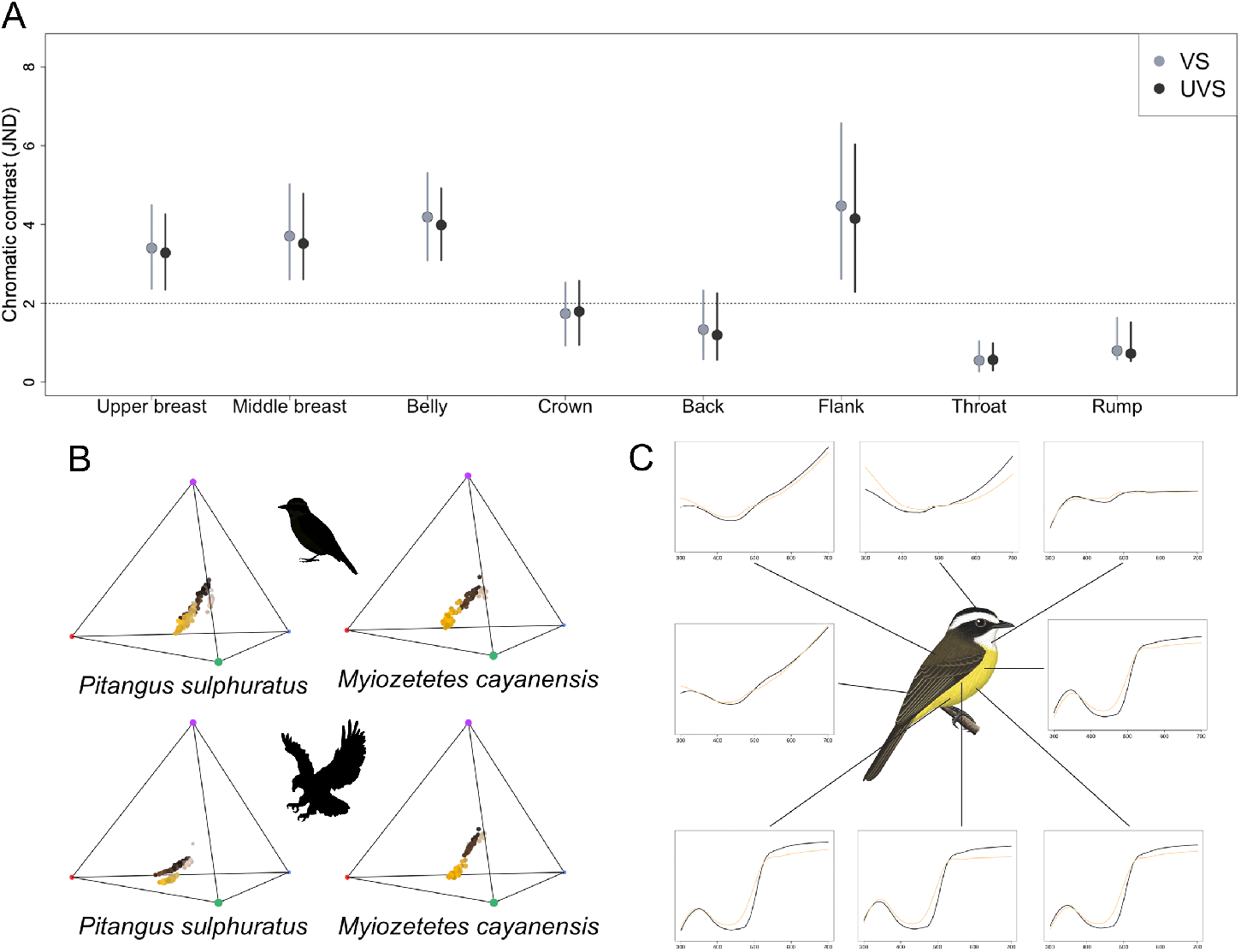
A hypothetical pair of model (*Pitangus sulphuratus*) and mimic (*Myiozetetes cayanensis*) species of flycatchers which we found are distinguishable in plumage coloration under visual models of discrimination abilities of putative competitors (UVS, passerines) and predators (VS, raptors). **A)** Comparison of color distances (in units of chromatic contrast or just noticeable differences JNDs) by patch given the UVS (black) and the VS (gray) vision models. The dotted horizontal line indicates JND=2, below which the pair of colors is considered to be indistinguishable by birds. Points and bars indicate the bootstrapped mean value and 95% CI’s respectively. **B)** Distribution of the color volume of each species in the tetrahedral color space using UVS and VS models. Volumes occupied by individual species differ between vision models; for a given vision model, color spaces occupied by each species are distinct. **C)** Reflectance curves for each plumage patch corrected by mean brilliance, with curve colors representing each of the two species being compared. There is a marked difference between model and mimic species ventral reflectance curves in regards to hue around 500nm. Illustration by Ayerbe-Quiñones [36] reproduced with permission from the author.

**Supplementary Figure 6.**
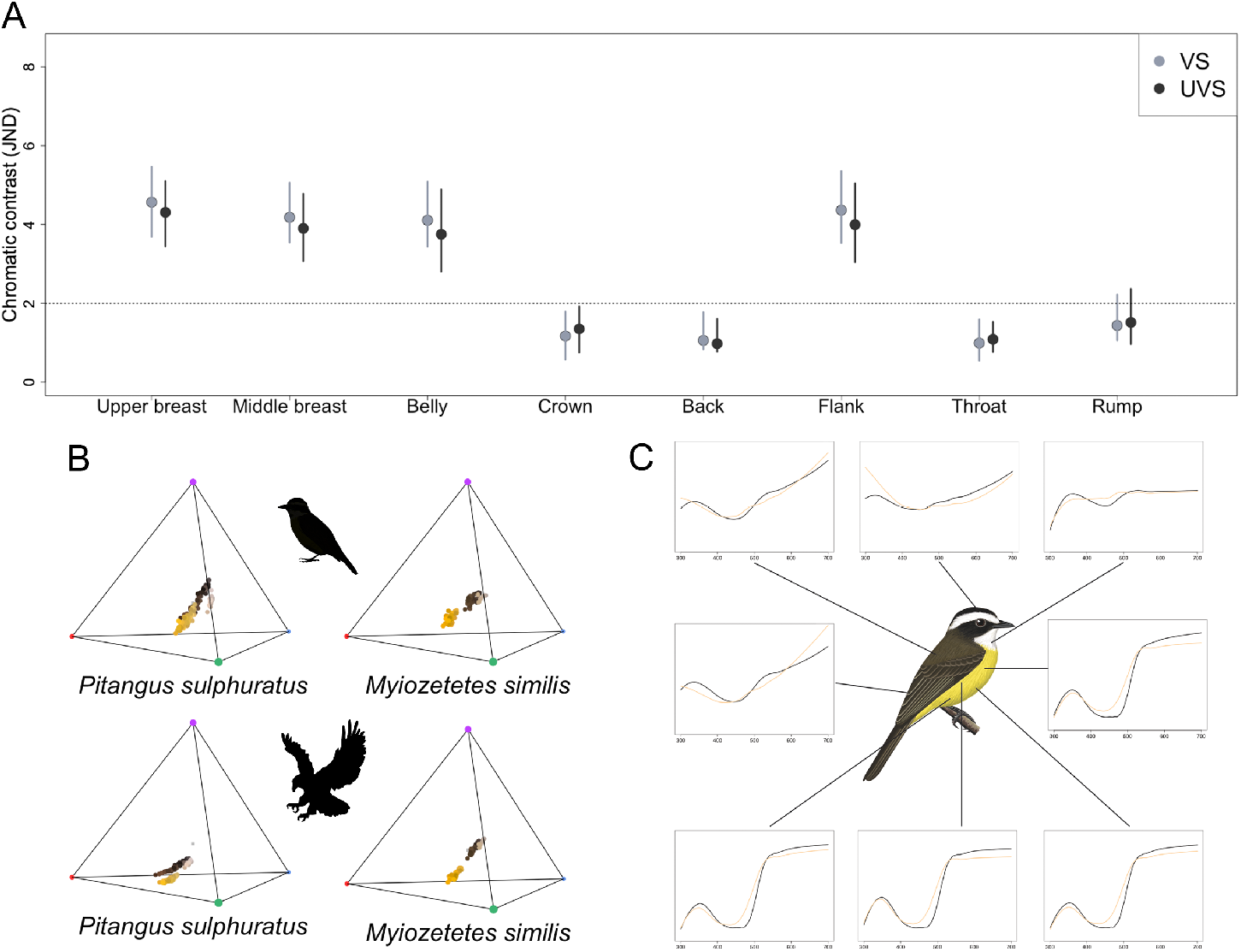
A hypothetical pair of model (*Pitangus sulphuratus*) and mimic (*Myiozetetes similis*) species of flycatchers which we found are distinguishable in plumage coloration under visual models of discrimination abilities of putative competitors (UVS, passerines) and predators (VS, raptors). **A)** Comparison of color distances (in units of chromatic contrast or just noticeable differences JNDs) by patch given the UVS (black) and the VS (gray) vision models. The dotted horizontal line indicates JND=2, below which the pair of colors is considered to be indistinguishable by birds. Points and bars indicate the bootstrapped mean value and 95% CI’s respectively. **B)** Distribution of the color volume of each species in the tetrahedral color space using UVS and VS models. Volumes occupied by individual species differ between vision models; for a given vision model, color spaces occupied by each species are distinct. **C)** Reflectance curves for each plumage patch corrected by mean brilliance, with curve colors representing each of the two species being compared. There is a marked difference between model and mimic species ventral reflectance curves in regards to hue around 500nm. Illustration by Ayerbe-Quiñones [36] reproduced with permission from the author.

**Supplementary Figure 7.**
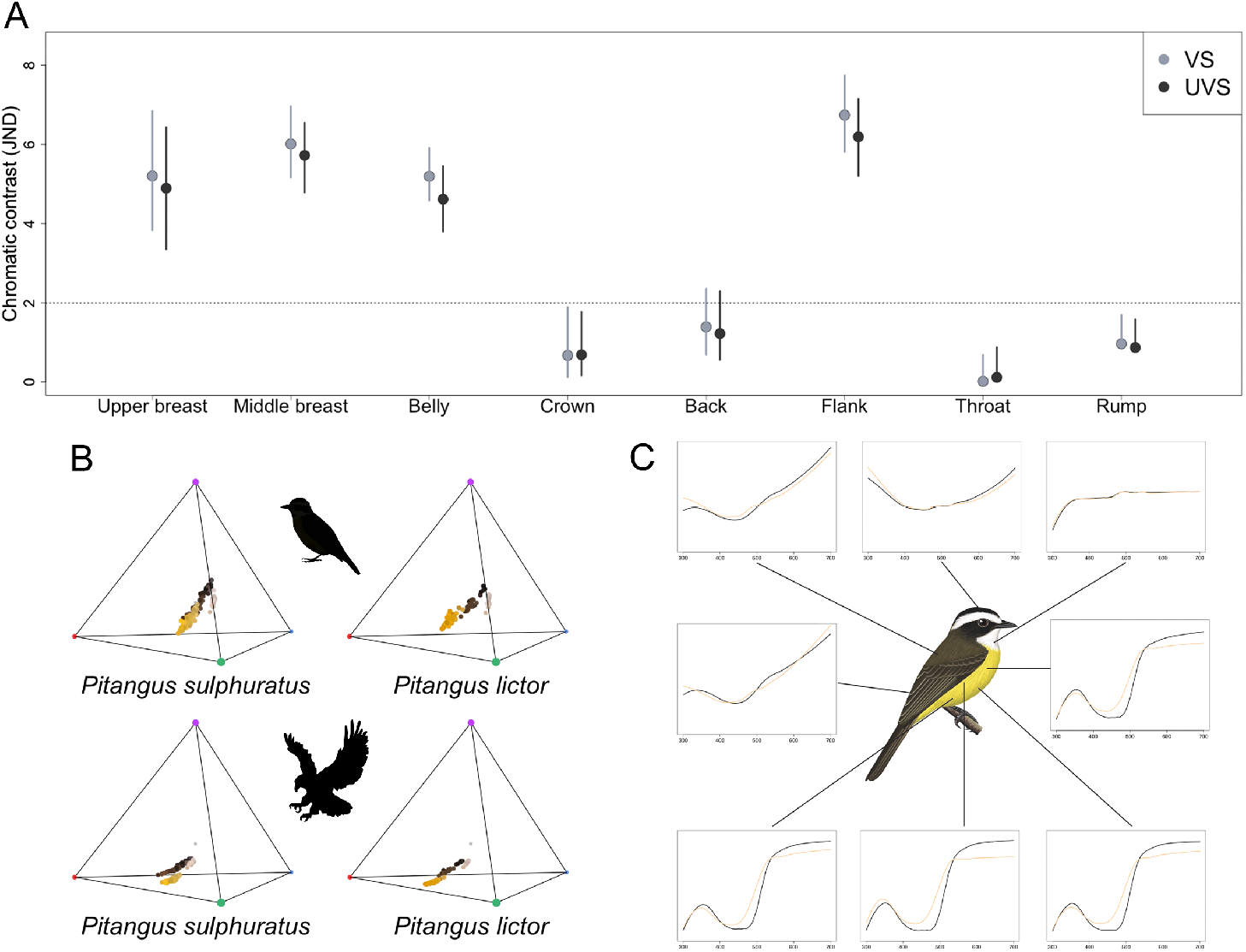
A hypothetical pair of model (*Pitangus sulphuratus*) and mimic (*Pitangus lictor*) species of flycatchers which we found are distinguishable in plumage coloration under visual models of discrimination abilities of putative competitors (UVS, passerines) and predators (VS, raptors). **A)** Comparison of color distances (in units of chromatic contrast or just noticeable differences JNDs) by patch given the UVS (black) and the VS (gray) vision models. The dotted horizontal line indicates JND=2, below which the pair of colors is considered to be indistinguishable by birds. Points and bars indicate the bootstrapped mean value and 95% CI’s respectively. **B)** Distribution of the color volume of each species in the tetrahedral color space using UVS and VS models. Volumes occupied by individual species differ between vision models; for a given vision model, color spaces occupied by each species are distinct. **C)** Reflectance curves for each plumage patch corrected by mean brilliance, with curve colors representing each of the two species being compared. There is a marked difference between model and mimic species ventral reflectance curves in regards to hue around 500nm. Illustration by Ayerbe-Quiñones (2018) reproduced with permission from the author.

**Supplementary Table 4.** Total and relative volume for all species using the UVS and VS models.

|  | UVS |  |  |  |  |  |
| --- | --- | --- | --- | --- | --- | --- |
|  | *M. pitangua* | *M. cayanensis* | *M. similis* | *P. inornata* | *P. lictor* | *P. sulphuratus* |
| Total volume | 0.00058 | 0.00084 | 0.00042 | 0.00056 | 0.00080 | 0.00109 |
| Relative volume | 0.00266 | 0.00393 | 0.00194 | 0.00259 | 0.00367 | 0.00503 |
|  | VS |  |  |  |  |  |
|  | *M. pitangua* | *M. cayanensis* | *M. similis* | *P. inornata* | *P. lictor* | *P. sulphuratus* |
| Total volume | 0.00026 | 0.00039 | 0.00019 | 0.00024 | 0.00035 | 0.00053 |
| Relative volume | 0.00125 | 0.00179 | 0.00088 | 0.00112 | 0.00166 | 0.00246 |

**Supplementary Table 5.** Centroid values of underpart patches for all species using the UVS and VS models.

| Upper Breast |  |  |  |  |  |  |  |  |  |  |  |  |
| --- | --- | --- | --- | --- | --- | --- | --- | --- | --- | --- | --- | --- |
|  | UVS |  |  |  |  |  | VS |  |  |  |  |  |
|  | *M. pitangua* | *M. cayanensis* | *M. similis* | *P. inornata* | *P. lictor* | *P. sulphuratus* | *M. pitangua* | *M. cayanensis* | *M. similis* | *P. inornata* | *P. lictor* | *P. sulphuratus* |
| u centroid | 0.14033 | 0.13016 | 0.13410 | 0.12556 | 0.13642 | 0.12876 | 0.08573 | 0.08090 | 0.08397 | 0.08570 | 0.08654 | 0.09722 |
| s centroid | 0.07265 | 0.07783 | 0.06695 | 0.07238 | 0.06354 | 0.12285 | 0.11865 | 0.12706 | 0.10920 | 0.11294 | 0.10250 | 0.17621 |
| m centroid | 0.37774 | 0.37909 | 0.37540 | 0.37484 | 0.37166 | 0.36446 | 0.38203 | 0.37936 | 0.37941 | 0.37488 | 0.37705 | 0.35408 |
| l centroid | 0.40926 | 0.41291 | 0.42352 | 0.42722 | 0.42836 | 0.38393 | 0.41356 | 0.41269 | 0.42740 | 0.42644 | 0.43391 | 0.37253 |
| Middle breast |  |  |  |  |  |  |  |  |  |  |  |  |
|  | UVS |  |  |  |  |  | VS |  |  |  |  |  |
|  | *M. pitangua* | *M. cayanensis* | *M. similis* | *P. inornata* | *P. lictor* | *P. sulphuratus* | *M. pitangua* | *M. cayanensis* | *M. similis* | *P. inornata* | *P. lictor* | *P. sulphuratus* |
| u centroid | 0.13994 | 0.11624 | 0.12870 | 0.12137 | 0.12151 | 0.11452 | 0.08996 | 0.07394 | 0.08335 | 0.07633 | 0.07318 | 0.08572 |
| s centroid | 0.08001 | 0.07367 | 0.06924 | 0.06119 | 0.05308 | 0.11946 | 0.12644 | 0.12139 | 0.11219 | 0.10252 | 0.09210 | 0.17597 |
| m centroid | 0.37574 | 0.38461 | 0.37979 | 0.38105 | 0.38387 | 0.37097 | 0.37762 | 0.38240 | 0.38115 | 0.38316 | 0.38837 | 0.35779 |
| l centroid | 0.40429 | 0.42546 | 0.42226 | 0.43635 | 0.44155 | 0.39504 | 0.40596 | 0.42225 | 0.42329 | 0.43799 | 0.44633 | 0.38052 |
| Belly |  |  |  |  |  |  |  |  |  |  |  |  |
|  | UVS |  |  |  |  |  | VS |  |  |  |  |  |
|  | *M. pitangua* | *M. cayanensis* | *M. similis* | *P. inornata* | *P. lictor* | *P. sulphuratus* | *M. pitangua* | *M. cayanensis* | *M. similis* | *P. inornata* | *P. lictor* | *P. sulphuratus* |
| u centroid | 0.10875 | 0.10101 | 0.12559 | 0.12893 | 0.12142 | 0.10940 | 0.06734 | 0.06270 | 0.08140 | 0.08910 | 0.07739 | 0.07915 |
| s centroid | 0.06419 | 0.06632 | 0.06883 | 0.07521 | 0.05991 | 0.11538 | 0.10901 | 0.11497 | 0.11209 | 0.11355 | 0.09860 | 0.17425 |
| m centroid | 0.39160 | 0.39476 | 0.38168 | 0.36843 | 0.37891 | 0.37466 | 0.39020 | 0.39008 | 0.38240 | 0.36952 | 0.38166 | 0.36110 |
| l centroid | 0.43544 | 0.43789 | 0.42388 | 0.42741 | 0.43972 | 0.40056 | 0.43342 | 0.43223 | 0.42409 | 0.42781 | 0.44235 | 0.38548 |
| Flank |  |  |  |  |  |  |  |  |  |  |  |  |
|  | UVS |  |  |  |  |  | VS |  |  |  |  |  |
|  | *M. pitangua* | *M. cayanensis* | *M. similis* | *P. inornata* | *P. lictor* | *P. sulphuratus* | *M. pitangua* | *M. cayanensis* | *M. similis* | *P. inornata* | *P. lictor* | *P. sulphuratus* |
| u centroid | 0.14215 | 0.12342 | 0.14417 | 0.13942 | 0.12169 | 0.13744 | 0.09236 | 0.07307 | 0.08846 | 0.08780 | 0.07200 | 0.09371 |
| s centroid | 0.08074 | 0.06985 | 0.06887 | 0.06597 | 0.05082 | 0.11966 | 0.12626 | 0.11822 | 0.11323 | 0.10765 | 0.08841 | 0.17975 |
| m centroid | 0.37521 | 0.38475 | 0.37650 | 0.37462 | 0.38166 | 0.3662 | 0.37743 | 0.38590 | 0.38211 | 0.37958 | 0.38745 | 0.35825 |
| l centroid | 0.40188 | 0.42196 | 0.41045 | 0.41999 | 0.44581 | 0.37670 | 0.40394 | 0.42279 | 0.41618 | 0.42498 | 0.45211 | 0.36826 |

## References

1. Losos JB. 2011 Convergence, adaptation, and constraint. Evolution 65, 1827–40. (doi: 10.1111/j.1558-5646.2011.01289.x)

2. Endler JA. 1986 Natural Selection in the Wild. Princeton University Press.

3. Lopes LE, Chaves AV, de Aquino MM, Silveira LF, dos Santos FR. 2018 The striking polyphyly of Suiriri: Convergent evolution and social mimicry in two cryptic Neotropical birds. J. Zool. Syst. Evol. Res. 56, 270–279. (doi: 10.1111/jzs.12200)

4. Price T, Pavelka M. 1996 Evolution selection of a colour pattern: history, development and selection. J. Evol. Biol. 9, 451–470.

5. Brakefield PM. 2006 Evo-devo and constraints on selection. Trends Ecol. Evol. 21, 362–368. (doi: 10.1016/j.tree.2006.05.001)

6. McGhee G. 2012 Convergent Evolution: Limited forms most beautiful. Evol. Dev. 14, 311–312. (doi: 10.1111/j.1525-142X.2012.00547.x)

7. Prum RO. 2014 Interspecific social dominance mimicry in birds. Zool. J. Linn. Soc. 172, 910–941. (doi: 10.1111/zoj.12192)

8. Jønsson KA, Delhey K, Sangster G, Ericson PGP, Irestedt M. 2016 The evolution of mimicry of friarbirds by orioles (Aves: Passeriformes) in Australo-Pacific archipelagos. Proc. R. Soc. B Biol. Sci. 283, 20160409. (doi: 10.1098/rspb.2016.0409)

9. Stoddard MC. 2012 Mimicry and masquerade from the avian visual perspective. Curr. Zool. 58, 630–648. (doi: 10.1093/czoolo/58.4.630)

10. Laiolo P. 2017 Phenotypic similarity in sympatric crow species: Evidence of social convergence? Evolution (N. Y). 71, 1051–1060. (doi: 10.1111/evo.13195)

11. Cody ML, Brown JH. 1970 Character Convergence in Mexican Finches. Evolution (N. Y). 24, 304. (doi: 10.2307/2406806)

12. Davies N, Welbergen J. 2008 Cuckoo-hawk mimicry? An experimental test. Proc. R. Soc. B Biol. Sci. 275, 1817–1822. (doi: 10.1098/rspb.2008.0331)

13. Leighton GM, Lees AC, Miller ET. 2018 The hairy-downy game revisited: an empirical test of the interspecific social dominance mimicry hypothesis. Anim. Behav. 137, 141–148. (doi: 10.1016/j.anbehav.2018.01.012)

14. Losos JB. 1998 Contingency and Determinism in Replicated Adaptive Radiations of Island Lizards. Science (80-.). 279, 2115–2118. (doi: 10.1126/science.279.5359.2115)

15. Miller ET, Leighton GM, Freeman BG, Lees AC, Ligon RA. 2019 Ecological and geographical overlap drive plumage evolution and mimicry in woodpeckers. Nat. Commun. 10, 1602. (doi: 10.1038/s41467-019-09721-w)

16. Barnard CJ. 1979 Predation and the Evolution of Social Mimicry in Birds. Am. Nat. 113, 613–618. (doi: 10.1086/283419)

17. Barnard CJ. 1982 Social Mimicry and Interspecific Exploitation. Am. Nat. 120, 411–415. (doi: 10.1086/284000)

18. Moynihan M. 1968 Social mimicry: character convergence versus character displacement. Evolution (N. Y). 22, 315–331. (doi: 10.2307/2406531)

19. Prum RO, Samuelson L. 2012 The Hairy-Downy Game: A model of interspecific social dominance mimicry. J. Theor. Biol. 313, 42–60. (doi: 10.1016/j.jtbi.2012.07.019)

20. Diamond J. 1982 Mimicry of Friarbirds by Orioles. Auk 99, 187–196.

21. Prum RO, Samuelson L. 2016 Mimicry Cycles, Traps, and Chains: The Coevolution of Toucan and Kiskadee Mimicry. Am. Nat. 187, 753–764. (doi: 10.1086/686093)

22. Dumbacher JP, Fleischer RC. 2001 Phylogenetic evidence for colour pattern convergence in toxic pitohuis: Müllerian mimicry in birds? Proc. R. Soc. London. Ser. B Biol. Sci. 268, 1971–1976. (doi:10.1098/rspb.2001.1717)

23. Gomez D, Théry M. 2007 Simultaneous Crypsis and Conspicuousness in Color Patterns: Comparative Analysis of a Neotropical Rainforest Bird Community. Am. Nat. 169, S42–S61. (doi: 10.1086/510138)

24. Mueller HC. 1971 Oddity and specific searching image more important than conspicuousness in prey selection. Nature 233, 345–346. (doi: 10.1038/233345a0)

25. Cuthill IC, Partridge JC, Bennett ATD, Church SC, Hart NS, Hunt S. 2000 Ultraviolet Vision in Birds. Adv. Study Behav. 29, 159–214. (doi: 10.1016/S0065-3454(08)60105-9)

26. Håstad O, Victorsson J, Ödeen A. 2005 Differences in color vision make passerines less conspicuous in the eyes of their predators. Proc. Natl. Acad. Sci. U. S. A. 102, 6391–6394. (doi:10.1073/pnas.0409228102)

27. Acosta-Chaves V, Granados F, Araya D. 2012 Predation of Long-tailed Silky Flycatcher (*Ptilogonys caudatus*) by Ornate Hawk-Eagle (*Spizaetus ornatus*) in a cloud forest of Costa Rica. Rev. Bras. Ornitol. 20, 451–452.

28. Amar A, Thirgood S, Pearce-Higgins J, Redpath S. 2008 The impact of raptors on the abundance of upland passerines and waders. Oikos 117, 1143–1152. (doi: 10.1111/j.0030-1299.2008.16769.x)

29. Thomson RL, Tomás G, Forsman JT, Broggi J, Mönkkönen M. 2010 Predator proximity as a stressor in breeding flycatchers: Mass loss, stress protein induction, and elevated provisioning. Ecology 91, 1832–1840. (doi: 10.1890/09-0989.1)

30. Gotmark F. 1995 Black-and-white plumage in male pied flycatchers (*Ficedula hypoleuca*) reduces the risk of predation from sparrowhawks (*Accipiter nisus*) during the breeding season. Behav. Ecol. 6, 22–26. (doi: 10.1093/beheco/6.1.22)

31. Slagsvold T, Dale S, Kruszewicz A. 1995 Predation favours cryptic coloration in breeding male pied flycatchers. Anim. Behav. 50, 1109–1121. (doi: 10.1016/0003-3472(95)80110-3)

32. Mueller HC. 1975 Hawks Select Odd Prey. Science (80-.). 188, 953–954. (doi: 10.1126/science.188.4191.953)

33. Endler JA. 1978 A predator’s view of animal color patterns. In Evolutionary biology, pp. 319–364. Springer.

34. Dunning JB. 2007 CRC Handbook of Avian Body Masses. CRC Press. (doi: 10.1201/9781420064452)

35. Hilty SL, Brown B. 1986 A guide to the birds of Colombia. Princenton University Press.

36. Ayerbe-Quiñones F. 2018 Guía ilustrada de la avifauna colombiana. Wildlife Conservation Society (WCS).

37. Fitzpatrick J et al. 2004 Volume 9, cotingas to P. In Handbook of the Birds of the World - Volume 9 Cotingas to Pipits and Wagtails (eds J del Hoyo, A Eliott, DA Christie), Barcelona: Lynx Edicions.

38. Maia R, Eliason CM, Bitton PP, Doucet SM, Shawkey MD. 2013 pavo: An R package for the analysis, visualization and organization of spectral data. Methods Ecol. Evol. 4, 906–913. (doi: 10.1111/2041-210X.12069)

39. Armenta JK, Dunn PO, Whittingham LA. 2008 Effects of specimen age on plumage color. Auk 125, 803–808. (doi: 10.1525/auk.2008.07006)

40. Maia R, White TE. 2018 Comparing colors using visual models. Behav. Ecol. 29, 649–659. (doi:10.1093/beheco/ary017)

41. Maia R, Gruson H, Endler JA, White TE. 2019 pavo 2: New tools for the spectral and spatial analysis of colour in r. Methods Ecol. Evol. 10, 1097–1107. (doi: 10.1111/2041-210X.13174)

42. Vorobyev M, Osorio D. 1998 Receptor noise as a determinant of colour thresholds. Proc. R. Soc. B Biol. Sci. 265, 351–358. (doi: 10.1098/rspb.1998.0302)

43. Vorobyev M, Brandt R, Peitsch D, Laughlin SB, Menzel R. 2001 Colour thresholds and receptor noise: Behaviour and physiology compared. Vision Res. 41, 639–653. (doi: 10.1016/S0042-6989(00)00288-1)

44. Vorobyev, Osorio D, Bennett ATD, Marshall NJ, Cuthill IC. 1998 Tetrachromacy, oil droplets and bird plumage colours. J. Comp. Physiol. A Sensory, Neural, Behav. Physiol. 183, 621–633. (doi: 10.1007/s003590050286)

45. Endler JA. 1993 The Color of Light in Forests and Its Implications. Ecol. Monogr. 63, 1–27. (doi: 10.2307/2937121)

46. Maia R, White T, Eliason C, Bitton PP. 2017 Package ‘pavo’. Packag. ‘pavo’. See https://cran.r-project.org/web/packages/pavo/pavo.pdf.

47. Oksanen J, Kindt R, Legendre P, O’Hara B, Simpson GL, Solymos PM, Stevens MHH, & Wagner H. 2008 The vegan package. Community Ecol. Packag., 190. (doi: 10.4135/9781412971874.n145)

48. Siddiqi A. 2004 Interspecific and intraspecific views of color signals in the strawberry poison frog *Dendrobates pumilio*. J. Exp. Biol. 207, 2471–2485. (doi: 10.1242/jeb.01047)

49. Bitton P-P, Janisse K, Doucet SM. 2017 Assessing Sexual Dicromatism: The Importance of Proper Parameterization in Tetrachromatic Visual Models. PLoS One 12, e0169810. (doi: 10.1371/journal.pone.0169810)

50. Hart NS. 2001 The Visual Ecology of Avian Photoreceptors. Prog. Retin. Eye Res. 20, 675–703. (doi: 10.1016/S1350-9462(01)00009-X)

51. Stoddard MC, Prum RO. 2008 Evolution of Avian Plumage Color in a Tetrahedral Color Space: A Phylogenetic Analysis of New World Buntings. Am. Nat. 171, 755–776. (doi: 10.1086/587526)

52. Endler JA, Mielke PW. 2005 Comparing entire colour patterns as birds see them. Biol. J. Linn. Soc. 86, 405–431. (doi: 10.1111/j.1095-8312.2005.00540.x)

53. Dalbosco Dell’Aglio D, Troscianko J, McMillan WO, Stevens M, Jiggins CD. 2018 The appearance of mimetic Heliconius butterflies to predators and conspecifics. Evolution (N. Y). 72, 2156–2166. (doi: 10.1111/evo.13583)

54. Müller F. 1879 Ituna and Thyridia: a remarkable case of mimicry in butterflies. Proc. Entomol. Soc. London 1879, 20–29.

55. Garg KM, Sam K, Chattopadhyay B, Sadanandan KR, Koane B, Ericson PGP, Rheindt FE. 2019 Gene Flow in the Müllerian Mimicry Ring of a Poisonous Papuan Songbird Clade (Pitohui; Aves). Genome Biol. Evol. 11, 2332–2343. (doi: 10.1093/gbe/evz168)

56. Weckstein JD. 2005 Molecular phylogenetics of the Ramphastos toucans: implications for the evolution of morphology, vocalizations, and coloration. Auk 122, 1191–1209. (doi: 10.1642/0004-8038(2005)122[1191:MPOTRT]2.0.CO;2)

57. Motta-Junior JC. 2007 Ferruginous Pygmy-owl (*Glaucidium brasilianum*) predation on a mobbing Fork-tailed Flycatcher (*Tyrannus savana*) in south-east Brazil. Biota Neotrop. 7, 76–79. (doi: 10.1590/S1676-06032007000200038)

58. Selas V. 1993 Selection of avian prey by breeding sparrowhawks *Accipiter nisus* in southern Norway: the importance of size and foraging behaviour of prey. Ornis Fenn. 70, 144–154.

59. Clark WS, Wheeler BK. 2001 A Field Guide to Hawks of North America. Second. New York: Houghton Miffin Company.

60. Wallace AR. 1863 List of birds collected in the island of Bouru (one of the Moluccas), with descriptions of the new species. Proc. Zool. Soc. London, 18–36.

61. Wallace AR. 1869 The Malay Archipielago: The land of the orangutan, and the bird of paradise. A narrative of travel, with studies of man and nature. Macmillan and Co.

62. Schoener TW. 1983 Field Experiments on Interspecific Competition. Am. Nat. 122, 240–285. (doi: 10.1086/284133)

63. Fitzpatrick JW. 1980 Foraging behavior of neotropical tyrant flycatchers. Condor 82, 43–57. (doi: 10.2307/1366784)

64. Fitzpatrick JW. 1981 Search strategies of tyrant flycatchers. Anim. Behav. 29, 810–821. (doi: 10.1016/S0003-3472(81)80015-2)

65. Tyrrell LP, Teixeira LBC, Dubielzig RR, Pita D, Baumhardt P, Moore BA, Fernández-Juricic E. 2019 A novel cellular structure in the retina of insectivorous birds. Sci. Rep. 9, 15230. (doi: 10.1038/s41598-019-51774-w)

66. Ensminger AL, Fernández-Juricic E. 2014 Individual variation in cone photoreceptor density in house sparrows: Implications for between-individual differences in visual resolution and chromatic contrast. PLoS One (doi: 10.1371/journal.pone.0111854)

